# Modeling Risk Group 4 virus infection and antiviral treatment in microfluidic lung organ-on-chips in maximum containment laboratories

**DOI:** 10.64898/2026.08.24.745299

**Authors:** Sushma M. Bhosle, Julie P. Tran, Shuiqing Yu, Jillian Geiger, Arpita Das, Scott M. Anthony, Bapi Pahar, Rebecca Bernbaum-Cutler, Deja F. P. Rivera, Ian Crozier, Jiro Wada, Anya Crane, Gustavo Palacios, Nicole C. Kleinstreuer, Jens H. Kuhn, Gabriella Worwa

## Abstract

Development of candidate countermeasures against human pathogens frequently includes nonhuman animal experimentation. Preclinical animal pathogen exposure studies are conducted to model diseases and accumulate preliminary and hypothetically translatable data to inform and justify the design of clinical trial evaluation of countermeasure safety and efficacy. In addition to frequent ethical critiques, challenges associated with animal experimentation include considerable resources needed to achieve statistical power and robustness, replicability and reproducibility concerns, potentially compromised objectivity through lack of blinding, fundamental species-specific biological differences, and risk of unpredictable pathogen adaptation to the experimental animal. Recent U.S. and U.K. government initiatives aim to reduce animal experimentation by complementing or potentially replacing them with new approach methodologies (NAMs), i.e., increasingly sophisticated *in silico*, *in chemico*, and *in vitro* approaches. We piloted development of one type of NAM, organ-on-chips (OOCs), in the highly challenging environment of a maximum (biosafety level 4) containment laboratory. Using a Risk Group 4 virus, Nipah virus (NiV), and two types of lung OOCs seeded with human or porcine cells, we demonstrated the recapitulation of key features of NiV lung infection, including viral infection, replication, and translocation, that are associated with proinflammatory cytokine secretion, immune cell recruitment, and disruption of the air–liquid interface barrier. We reproduced the known anti-NiV activity of remdesivir and evaluated that of another potential antiviral, zotatifin. Our results pave the way for similar applications of advanced microphysiological systems for modeling infections caused by high-consequence viruses.

## INTRODUCTION

Animal experimentation has been a major component of preclinical research and development, including to inform subsequent clinical trial evaluation that is required for licensure of medical countermeasures (MCMs) for the diagnosis, treatment, or prevention of human high-consequence infections. Briefly, nonhuman animals identified as susceptible to a particular human pathogen are hypothesized as translational proxies for humans, i.e., nonhuman animals that are best suited to model a particular aspect of a human infection and/or disease after exposure to the pathogen in absence of, before, during, or after administration of candidate MCMs. Pathophysiological responses (the expression of disease) to the exposure and treatment are characterized in detail to project “translatability” to humans. If promising, preliminary data from animal models are sourced to justify, plan, and execute human clinical trials of MCMs, for instance during infectious disease outbreaks of novel or reemerging pathogens (<u>Korch et al.,</u> <u>2011</u>).

Historically, preclinical studies have not been required in all circumstances. For instance, “straight-to” human clinical trials could be directly performed if the safety profile of an MCM had already been established in humans and lack of efficacy was not expected to result in severe disease or the death of patients (e.g., in specific cases of nonlethal pathogens). However, with regard to “exotic” pathogens that emerge rarely and in unpredictable locations (making planning and execution of clinical trials logistically challenging) and/or for pathogens that cause severe morbidity and/or a high case-fatality rate (CFR; though controversially, querying whether the inclusion of an untreated/placebo group in a clinical trial is ethical), animal experimentation has been largely considered an absolute requirement. In the United States, specifically designed legislation, colloquially known as the U.S. Food and Drug Administration (FDA) “Animal Rule”, outlined and all but required a threshold of animal model data to permit direct administration of candidate MCMs, either under expanded-access or in clinical trial, to patients in times of emergency (<u>Food and Drug Administration, 2024</u>).

However, animal experimentation has also been frequently criticized for ethical reasons; the considerable financial and logistical resource requirements; challenges with achieving statistical power; a lack of robustness and objectivity; concerns about replicability and reproducibility; translational relevance across species; and concerns about unwanted pathogen adaptation. Adding concern is the accumulated experience that only rarely has this pathway actually resulted in new licensed MCMs (<u>Greek & Menache, 2013</u>; <u>Lin, 1995</u>). Finally, rigorous and well-executed clinical trials of MCMs have been able to be conducted in humans in infectious disease outbreak settings previously thought intractable to these efforts; furthermore, results from these trials have challenged previous assumptions about the translatability of MCM efficacy from animal models to humans.

Recent technological and methodological progress is challenging the notion that classical animal model experiments cannot be complemented or even potentially replaced by experiments utilizing new approach methodologies (NAMs), including perfused human organs, precision-cut tissue slices, two-dimensional (2D) hydrogels, three-dimensional (3D) bioprinted materials, spheroids and organoids, organ-on-chips (OOCs), and *in silico* approaches (e.g., digital twins). NAMs are increasingly studied as human-biology-based approaches to recapitulate some aspects of *in vivo* microenvironments that are tractable for investigation of host–pathogen interactions, pathogen replication kinetics, and MCM activity, while overcoming inter-species (model animal vs. human) differences (<u>Bhosle et al., 2018</u>; <u>de Melo et al., 2021</u>; <u>Dihan et al., 2024</u>; <u>Ghallab,</u> <u>2013</u>; <u>Ingber, 2003</u>; <u>Kitaeva et al., 2020</u>; <u>Ma et al., 2018</u>).

Consequently, NAMs became a focus in the FDA’s “Modernization Act 2.0” of 2022 (<u>U.S. Congress, 2022</u>), and the organization’s subsequent intention in 2025 to strategically phase out reliance on animal model data for preclinical MCM safety assessment, beginning with monoclonal antibodies (<u>Food and Drug Administration, 2025</u>). Also in 2025, the U.S. National Institutes of Health (NIH) announced its intention to prioritize human-based technologies for biomedical research (<u>National Institutes of Health, 2025b</u>).

Maximum (U.S.: biosafety level 4 [BSL-4]) containment laboratories focus on MCM research and development (R&D) against Risk Group 4 (RG-4) pathogens, i.e., “exotic” viruses associated with extraordinarily high CFRs in infected humans (<u>Richmond, 2002</u>; <u>U.S.</u> <u>Department of Health and Human Services et al., 2020</u>; <u>World Health Organization, 2017</u>). Typically, licensed MCMs are not identified or available for prevention or treatment of these infections; even when identified, conducting human clinical trials of safety and efficacy has been historically challenging in typical at-risk regions. The pace of MCM R&D has been limited by the fact that there are few maximum containment laboratories in operation due to maintenance costs, biosafety needs, and security requirements (<u>Richmond, 2002</u>; <u>World Health Organization,</u> <u>2017</u>). In the recent past, the FDA Animal Rule was the only path to licensure of MCMs targeting RG-4 pathogens, further limiting progress because only a subset of the maximum containment facilities could perform animal experimentation. Due to a recent NIH announcement that Notices of Funding Opportunities (NOFOs) exclusive to animal models would no longer be issued (<u>National Institutes of Health, 2025a</u>), NIH-supported maximum containment laboratories are motivated to incorporate NAMs into standard research portfolios.

With this in mind, we describe the setup, use, challenges, and solutions associated with experimental deployment of a particular NAM, i.e., OOCs, in a U.S. BSL-4 containment setting. OOCs are microfluidic devices consisting of channels seeded with primary cells in a pre- definable fashion, thus creating functional units that model the complex 3D anatomy and function of organs or tissues by emulating relevant microenvironmental, mechanical, and physiological forces (<u>Ingber, 2022</u>; <u>Tang et al., 2020</u>). For instance, a lung OOC contains a semipermeable membrane that separates two channels in which seeding of human donor primary microvascular endothelial and lung epithelial cells, respectively, architecturally model the lung microenvironment (<u>Huh et al., 2010</u>). Microengineered human lung OOCs have been used to model several noninfectious lung diseases (<u>Benam et al., 2016</u>; <u>Huh et al., 2012</u>; <u>Nawroth et al.,</u> <u>2020</u>) and in cancer and lung thrombosis models for drug screening evaluation of therapeutics (<u>Jain et al., 2018</u>; <u>Xu et al., 2013</u>). In virology, human lung OOCs have been developed to study of and screen antivirals against a scant few Risk Group 2 (i.e., influenza A virus) (<u>Koceva &</u> <u>Mosig, 2025</u>; <u>Si et al., 2021</u>) and Risk Group 3 viruses (i.e., severe acute respiratory syndrome coronavirus 2 [SARS-CoV-2]; <u>Man et al., 2025</u>; <u>Zhang et al., 2021</u>), but not yet against RG-4 viruses.

Here, as proof of principle, we outline the operational and experimental protocol to establish lung OOC capability in maximum containment; in a pilot study, via the use of human small airway and porcine alveolus lung OOCs exposed to an RG-4 virus, we subsequently describe methods to characterize viral infection, immune responses and immune cell recruitment, and infection-associated barrier disruption, including in the presence of both a recognized and a novel antiviral MCM. Broad application and adaptation of these methods in the BSL-4 setting supports the use of OOCs to study RG-4 viral infection and disease pathogenesis and to evaluate candidate MCMs in species-relevant NAMs.

## RESULTS

### Establishment and maintenance of air–liquid interfaces in human and porcine lung organ-on-chips

To establish initial organ-on-chip (OOC) capability in a maximum (biosafety level 4 [BSL-4]) containment setting, we strategically selected a commercially available system given the inherent initial advantages, including availability, material standardization and instruction, access to customer support, and (although still limited) the opportunity to capitalize on shared experiences and troubleshooting through recent literature or collaborative or community interest groups specific to microphysiological systems (MPS). For proof-of-principle setup, we selected lung OOCs that incorporated lung air–liquid interfaces (ALI; <u>Huh et al., 2010</u>) with the ultimate goal of emulating an infection with a Risk Group 4 (RG-4) pathogen that must be handled in BSL-4 containment (**Figure 1a**). We selected Nipah virus (NiV; *Mononegavirales*: *Henipavirus nipahense*) as our proof-of-principle RG-4 pathogen (<u>U.S. Department of Health and Human</u> <u>Services et al., 2020</u>) because its pathogenesis in accidental hosts (humans and domestic pigs [*Sus domesticus* Erxleben, 1777]) and experimental hosts (nonhuman primates, rodents) is generally well-characterized (<u>Spengler et al., 2025</u>) and because NiV has been designated a priority pathogen by the U.S. National Institute for Allergy and Infectious Diseases (<u>National</u> <u>Institute of Allergy and Infectious Diseases, 2024</u>) and other organizations (<u>World Health</u> <u>Organization, 2022</u>) due to the absence of licensed medical countermeasures (MCMs) for a severe disease with high CFRs.

**Figure 1.**
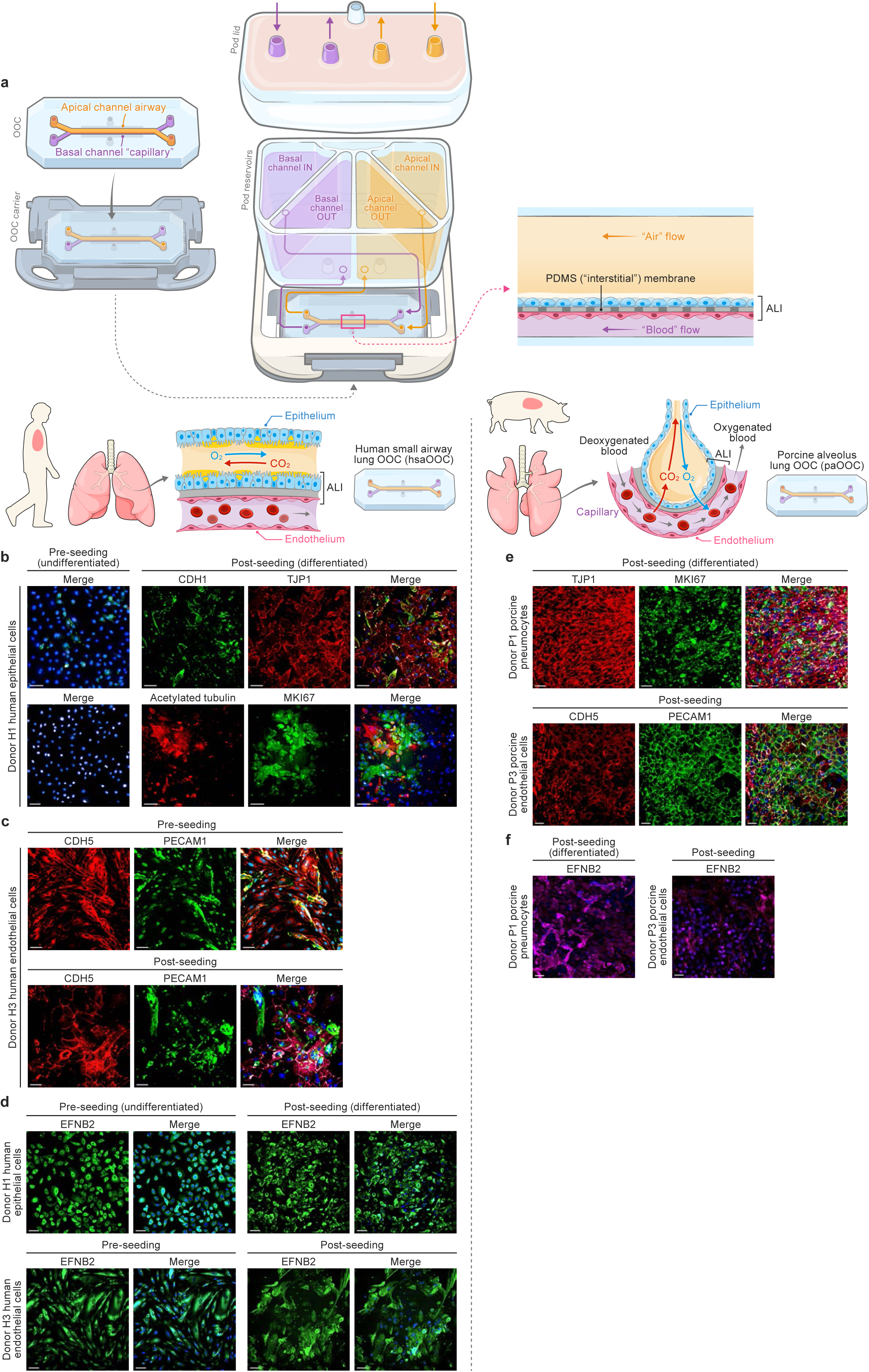
Establishment and maintenance of air–liquid interfaces in human and porcine lung organ-on-chips. **a**, Schematic of a generic lung organ-on-chip (OOC). Apical (“parenchymal”/“airway”/top; orange) and basal (“vascular”/“capillary”/bottom; purple) channels connected to their inlet/outlet pod reservoirs enable simultaneous “air” and “blood” flow. Differentiated donor primary epithelial (blue) and endothelial (red) cells cultured in the apical and basal channels of the OOC, respectively, cover a semi-permeable polydimethylsiloxane (PDMS) membrane, thereby simulating the lung interstitium and the air– liquid interface (ALI). **b–d,** Modelling a human small airway (drawings) on a specific lung OOC (hsaOOC). Immunostaining was performed on epithelial cells from human cell donors cultured in 96-well plates prior to differentiation (pre-seeding into OOCs) and differentiated cells after detaching them from hsaOOCs after 3 weeks under ALI conditions, as well as on endothelial cells from human cells pre- and post-seeding. Images were captured with a 20X objective and equal exposure and gain settings using a high-content imaging system. Shown are representative images from duplicate experiments for the human epithelial cell donor (Donor H1) and human endothelial cell donor (Donor H3). CO_2_, carbon dioxide; O_2_, oxygen. **b,** Immunofluorescence micrographs of epithelial cells stained with antibodies against characteristic tight-junction cadherin 1 (CDH1, green), tight junction protein 1 (TJP1, red) differentiation marker of proliferation Ki-67 (MKI67, green), and acetylated tubulin (red) markers. **c,** Immunofluorescence micrographs of human endothelial cells stained with antibodies against characteristic cadherin 5 (CDH5, red) and platelet and endothelial cell adhesion molecule 1 (PECAM1, green) markers. **d,** Immunofluorescence micrographs of epithelial and endothelial cells stained with antibodies against Nipah virus (NiV) host cell receptor ephrin B2 (EFNB2, green). **e–f,** Modelling a porcine alveolus lung OOC (paOOC; drawings). Shown are representative images from duplicate experiments for pneumocyte Donor P1 and endothelial cell Donor P3. **e,** Immunofluorescence micrographs of differentiated porcine pneumocytes and porcine endothelial cells in the apical and basal channels of the paOOCs, respectively. Epithelial cells were stained with antibodies against TJP1 (red) and MKI67 (green). Endothelial cells were stained with antibodies against CDH5 (red) and PECAM1 (green). **f**, Immunostaining of porcine pneumocytes and porcine endothelial cells stained with an antibody against NiV receptor EFNB2 (magenta). Nuclei in **b–f** were counterstained with Hoechst (blue). Scale bars, 100 µm.

BSL-4 laboratories are challenging workspaces that require highly trained personnel (<u>Richmond, 2002</u>; <u>U.S. Department of Health and Human Services et al., 2020</u>; <u>World Health</u> <u>Organization, 2017</u>). Beyond installation and maintenance of required equipment, establishing novel workflows and techniques in these facilities is difficult because safety protocols and other standard operating procedures (SOPs) need to be rigorously designed, continuously evaluated, and revised when necessary. Further, the personal protective equipment mandatory for BSL-4 work impacts staff dexterity and the time-intensive donning and doffing of safety suits make the initial setup cumbersome (<u>Janosko et al., 2016</u>; <u>Mazur et al., 2016</u>). In early development of this capacity, we strategically established mirror setups of OOC equipment and processes in a BSL-2 laboratory setting; this parallel system not only accelerated staff training and familiarization but also the development and troubleshooting of virus-free steps under much less stringent BSL-2 working conditions.

BSL-2 setup of a commercial MPS was relatively straightforward. To establish a human small airway lung OOC (hsaOOC; drawings on top of **Figures 1b–d**) we prepared both channels with manufacturer-proprietary activation and coating reagents, ensuring proper extracellular matrix (ECM) coating. We purchased frozen primary human small airway epithelial cells (from here on: human epithelial cells) sampled from two different human donors (Donor H1 and Donor H2) and frozen primary human lung microvascular endothelial cells (from here on: human endothelial cells) from one human donor (Donor H3; **Supplementary Table 1**). Cells were thawed and expanded in tissue culture flasks and then seeded into the basal (“vascular”/“capillary”/bottom) channels. After attachment of the human endothelial cells, the human epithelial cells were seeded into the apical (“parenchymal”/“airway”/top) channels (**Figure 1a**). After overnight cell settling and attachment, the seeded hsaOOCs were connected to the pod reservoirs and subsequently inserted into a culture module in which the flow rate of media was independently set to 45 µL/h through both hsaOOC channels. After confluence of cell layers was confirmed, media in the apical channels were replaced under the same flow rate to prepare the human epithelial cells for differentiation. Cell differentiation was induced by subsequent removal of the media and setting the flow rate to 0 µL/h (“air exposure”) in the apical channels while maintaining the 45 µL/h media flow rate in the basal channels. For follow-up experiments with hsaOOCs, the origin of human epithelial cells differed (from Donor H1 or Donor H2) but all endothelial cells originated from Donor H3 (→Hx/H3 hsaOOCs with x denoting the donor number). Human epithelial cell differentiation was evaluated by comparing human epithelial cells cultured in 96-well plates prior to hsaOOC seeding with cells cultured in the hsaOOCs after the ALI was established: characterization included immunohistochemical (IHC) staining with antibodies targeting tight-junction markers (cadherin 1 [CDH1] and tight-junction protein 1 [TJP1]) and differentiation markers (proliferation marker Ki-67 [MKI67] and acetylated tubulin). Fluorescence microscopy revealed that cells cultured in the 96-well plates, which did not undergo differentiation or establishment of ALI, lacked the expression of these markers. In contrast, well-defined marker expression was observed following establishment of ALI in hsaOOCs (example for Donor H1 shown in **Figure 1b**), indicating appropriate epithelial cell differentiation within hsaOOCs. Via a similar approach, we characterized human endothelial cells of the basal channels, including IHC with antibodies against endothelial cell-specific markers cadherin 5 (CDH5) and platelet and endothelial cell adhesion molecule 1 (PECAM1); we confirmed that marker expression in these cells was retained after hsaOOC seeding and establishment of the ALI (**Figure 1c)**. By IHC staining and microscopy, we also confirmed that ephrin B2 (EFNB2), the primary host cell receptor of NiV (<u>Bonaparte et al., 2005</u>; <u>Negrete et al.,</u> <u>2005</u>), was expressed on the human endothelial and human epithelial cells before and after seeding (**Figure 1d**).

As next proof-of-principle, we established the utility for nonhuman OOCs to enable comparative study of zoonotic viruses, including NiV, in disparate hosts. Because NiV was first discovered during a large epizootic among captive domestic pigs (*Sus domesticus* Erxleben, 1777), we targeted establishment of porcine OOCs. To demonstrate the versatility of the OOC system and to enable direct intra-species comparisons with this known zoonotic reservoir, we chose to emulate the domestic pig alveolus, i.e., to create porcine alveolus lung OOCs (from here on: paOOCs; drawings on top of **Figures 1e–f**). In establishing paOOCs, apical channels were seeded with porcine alveolar cells (from here on: porcine pneumocytes) originating from either of two porcine donors (Donor P1 or Donor P2) after the basal channels had been seeded with porcine microvascular endothelial cells (from here on: porcine endothelial cells) derived from a third porcine donor (Donor P3; **Supplementary Table 2**). Following the connection of the paOOCs to the pod reservoirs, we applied mechanical stretch via the culture module to mimic physiological forces experienced by tissues *in vivo*. Differentiation of porcine pneumocytes in the paOOCs was complete within 8 d after establishment of the ALI, and we confirmed appropriate porcine pneumocyte differentiation via staining with antibodies against differentiation markers TJP1 and MKI67. IHC staining was also used to verify expression of CDH5 and PECAM1 in porcine endothelial cells in the paOOCs. The NiV receptor EFNB2 was detected in both porcine endothelial cells and porcine pneumocytes (example for Donor P1 shown in **Figures 1e–f**).

Overall, we successfully established human small-airway-like and porcine alveolus-like lung OOCs with properly differentiated cells under ALI conditions in the absence (hsaOOCs) or presence (paOOCs) of mechanical stretch.

### Nipah virus disrupts the air–liquid interface barrier in human small airway lung organ-on-chips

In humans with severe disease, NiV infection presents with central nervous and respiratory system manifestations (<u>Spengler et al., 2025</u>). Significant differences in the clinical disease phenotype and severity associated with infections with Bangladesh (NiV-B) and Malaysia (NiV-M) isolates have been reported: although encephalitis is reported after both, NiV-B infection is associated with a higher frequency of respiratory disease and higher overall CFRs (<u>Clayton et al.,</u> <u>2012</u>; <u>Geisbert et al., 2010</u>; <u>Goldin et al., 2025</u>; <u>Gurley et al., 2007</u>; <u>Mire et al., 2016</u>; <u>Sazzad et</u> <u>al., 2013</u>; <u>Wong et al., 2002</u>). Consequently, we selected a NiV-B isolate for follow-up experiments.

After completion of cell differentiation (approximately 3 weeks after establishment of ALI), hsaOOCs were transferred from the BSL-2 to a BSL-4 laboratory; subsequently, the human epithelial cells (in the apical channels) were exposed to NiV at a multiplicity of infection (MOI) of 1 in medium for 2–2.5 h under flow and static conditions, mimicking a respiratory exposure. After incubation, the virus inocula were removed from the apical channels and the appropriate cell media were flowed through both channels at a constant rate. The standard workflow for exposure of OOCs to NiV, connection of OOCs to the culture module, and collection of assay samples is shown in **Figure 2a**. Effluents and channel cell lysates from OOCs were collected and analyzed by multiple assays (**Figure 2b**). First, we quantified NiV genome equivalents (GEq) by quantitative reverse transcription polymerase chain reaction (RT-qPCR) and evaluated NiV infectivity by plaque assay. In hsaOOCs, NiV GEq were detected in both apical and basal channels effluents collected 24 and 48 h after NiV exposure, indicating that NiV had transversed the polydimethylsiloxane (PDMS; “interstitial”) membranes to enter the basal channels soon after exposure. By RT-qPCR, at both time points, the apical channel effluents contained higher quantities of NiV GEq than the basal channel effluents, indicating higher replication at the site of exposure (**Figure 2c**). Similarly, plaque assay detected higher NiV titers in apical channel effluents at 24 and 48 h post-exposure than in basal channel effluents (**Figure 2d**). Together, RT-qPCR and plaque assay data suggest a delay of NiV translocation across the barrier. NiV GEq were also detected by RT-qPCR in lysates of both human epithelial and human endothelial cells and, unsurprisingly, cell lysate values were higher compared to the respective channels’ effluents (**Figure 2e**). Second, we detected syncytia formation, a hallmark of NiV infection (<u>Gamble et al., 2021</u>), by microscopy of the hsaOOC epithelial cells at 48 h (**Figure 2f**). hsaOOCs endothelial cells could not be examined by microscopy due to technical limitations in the BSL-4 laboratory. Third, we stained the hsaOOC cells with antibodies against NiV glycoprotein (G): NiV G expression was detected on human epithelial and human endothelial cell 48 h after NiV exposure (**Figure 2g**). Finally, we evaluated the integrity/permeability of the PDMS barrier between the epithelial and endothelial cell layers in the hsaOOCs with a barrier function test. The apparent permeability coefficient (P_app_) was quantified by introduction of fluorescent dye (Cascade Blue 3kDa) beads through the basal channels and measurement of the tracer amount in the effluent from both channels over time. In non-exposed hsaOOCs, the barrier permeability remained stable at 48 h but was considerably compromised after NiV infection (**Figure 2h**), suggesting that NiV first infected human epithelial cells, subsequently disrupted the ALI, and then transversed the “interstitial” membrane to infect human endothelial cells in the hsaOOCs.

**Figure 2.**
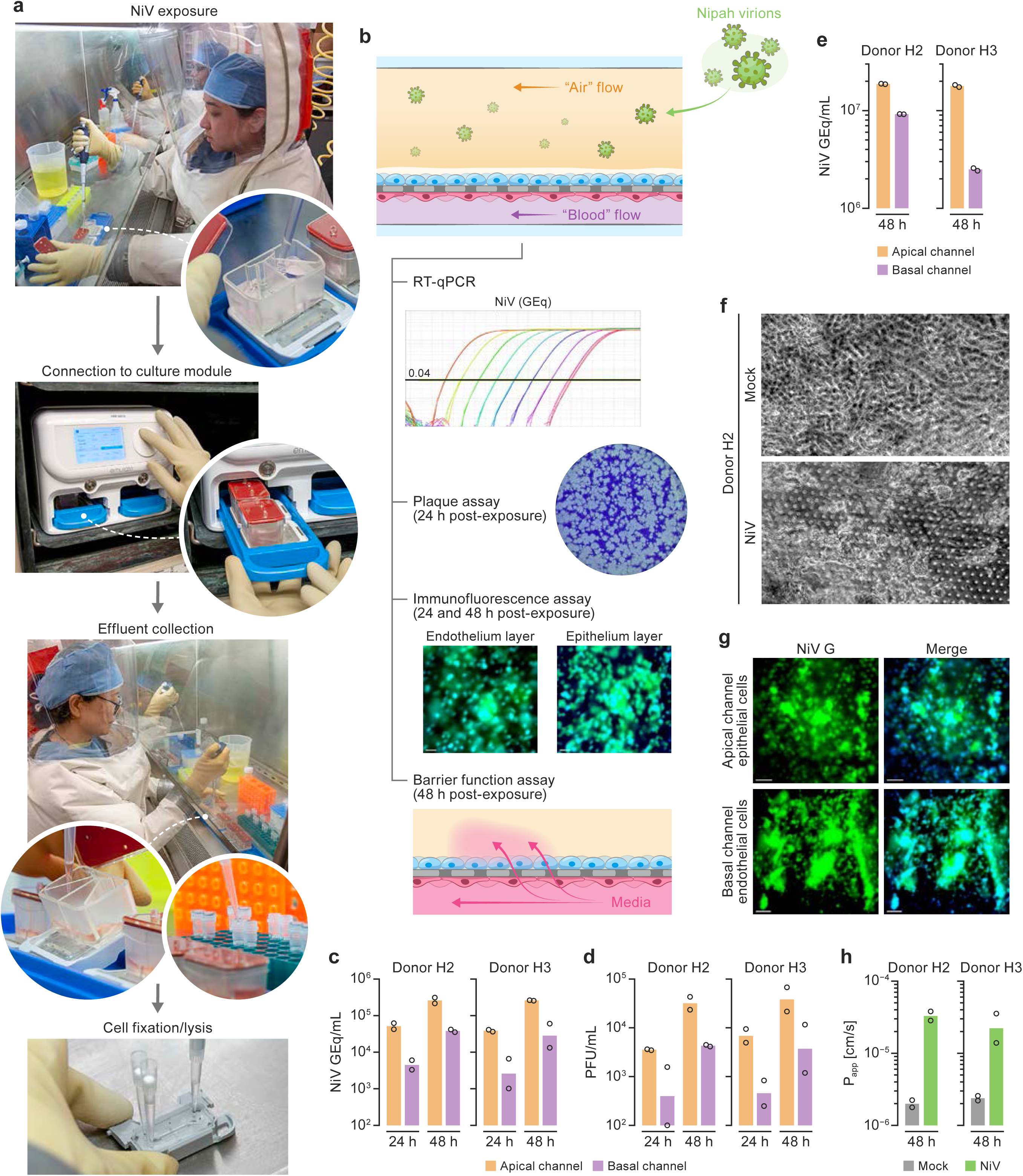
Nipah virus disrupts the air–liquid interface barrier in human small airway lung organ-on-chips. **a**, Workflow of Nipah virus (NiV) human small airway lung organ-on-chip (hsaOOC) exposure experiments. hsaOOCs were prepared using epithelial cells from either human Donor H1 or Donor H2 (apical channel) and endothelial cells from human Donor H3 (basal channel), yielding H1/H3 and H2/H3 hsaOOCs, in a biosafety level 2 (BSL-2) laboratory (not shown). Once cells were fully differentiated under air–liquid interface (ALI) conditions, transferred to a maximum containment (biosafety level 4) laboratory for exposure to Nipah virus (NiV) at 0 h via media infusion into the apical channel. hsaOOCs were then connected to the OOC system’s culture module for continuous media flow through both channels, followed by effluent collection and cell fixation/lysis/lysate collection at appropriate time points. Consent for publication of author photographs was obtained from all featured participants. **b,** Downstream assays performed on NiV-exposed hsaOOC effluents and cells included quantitative reverse transcription polymerase chain reaction (RT-qPCR) on hsaOOC effluents for detection of NiV genome equivalents (GEq), plaque assay titration for detection of infectious NiV, immunofluorescence assays for detection of cell-specific surface markers and the NiV host cell receptor, barrier function assays, proinflammatory cytokine quantification, and RT-qPCR arrays for quantification of human endothelial cell inflammation (not all assays are shown). **c,** NiV GEq quantified by RT-qPCR in effluents harvested from apical and basal channels after NiV exposure using duplicate hsaOOCs. Results are shown as the means of triplicate values with standard deviations (SDs). Assays were performed on samples from four human donors (*n*=4). **d,** Titers determined by plaque assay in hsaOOC effluents collected after NiV exposure (*n*=2). Data represent means of duplicates with SDs. **e,** NiV GEq quantified by RT-qPCR in cell lysates (*n*=2). Results are shown as the means of triplicate values with SDs. **f,** Syncytia formation visible by bright field microscopy in human epithelial cells at 48 h following exposure to mock-exposed control or NiV. **g,** Immunofluorescence assay reveals NiV glycoprotein G (green) in epithelial and endothelial cells at 48 h after NiV exposure. Shown are representative images for H1/H3 OOCs. Green: NiV G. Blue: cell nuclei (Hoechst staining). Images were captured with a 20X objective on a Leica microsystem; scale bar, 50 µm. **h,** Barrier permeability assessment after NiV exposure (hsaOOC; *n*=2). P_app_, apparent permeability coefficient. Results are shown as the means of coefficient values with SDs.

Together, these data demonstrate proof-of-principle feasibility for the use of hsaOOCs as an experimental platform to investigate the host–virus interaction for pneumotropic RG-4 pathogens in a BLS-4 setting.

### Small-molecule antivirals reduce Nipah virus titers in lung organ-on-chips

As further proof-of-principle, we determined if OOCs could be leveraged to evaluate the activity of known or novel small-molecule antiviral MCMs against RG-4 virus infections. We selected remdesivir, previously reported to be highly active against NiV infection *in vitro* and in experimentally exposed nonhuman primates (<u>Lo et al., 2019</u>) as a positive control. Based on promising results from two-dimensional (2D) cell culture drug screening assays (preliminary data, not shown) we also evaluated the anti-NiV activity of zotatifin, an inhibitor of eukaryotic translation initiation factor 4 alpha (EIF4A; <u>Gordon et al., 2020</u>; <u>Müller et al., 2021</u>; <u>Obermann</u> <u>et al., 2022</u>).

To evaluate the antiviral activity of these molecules, hsaOOCs were “treated” with the antivirals at the time of NiV exposure. Antiviral exposure was maintained in the hsaOOCs for 48 h after exposure by flowing media only (“untreated”), media + remdesivir (100 nM), or media + zotatifin (50 nM) continuously through both channels to mimic a therapeutic effect (**Figure 3a**). NiV GEq were significantly reduced in effluents from both channels at 24 and 48 h in hsaOOCs “treated” with either molecule compared to the “untreated” control (**Figures 3b–c**). hsaOOCs that received the antivirals maintained the ALI barrier, whereas permeability increased in the “untreated” controls (**Figure 3d**), suggesting a barrier-protective effects for both compounds. Consistent with RT-qPCR results, plaque assay data showed a significant reduction in NiV titers in the “treatment” groups compared to the “untreated” controls in apical and basal effluents (**Figures 3e–g**). Of interest, when compared to control hsaOOCs, NiV GEqs were considerably reduced in human epithelial and human endothelial cells in remdesivir-treated hsaOOCs, consistent with remdesivir’s mode of action as a nucleoside analog (<u>Lo et al., 2019</u>). However, in zotatifin-“treated” hsaOOCs, the reduction in NiV GEq was not as pronounced in the epithelial cells and varied in the endothelial cells among the different OOCs (**Figure 3h**).

**Figure 3.**
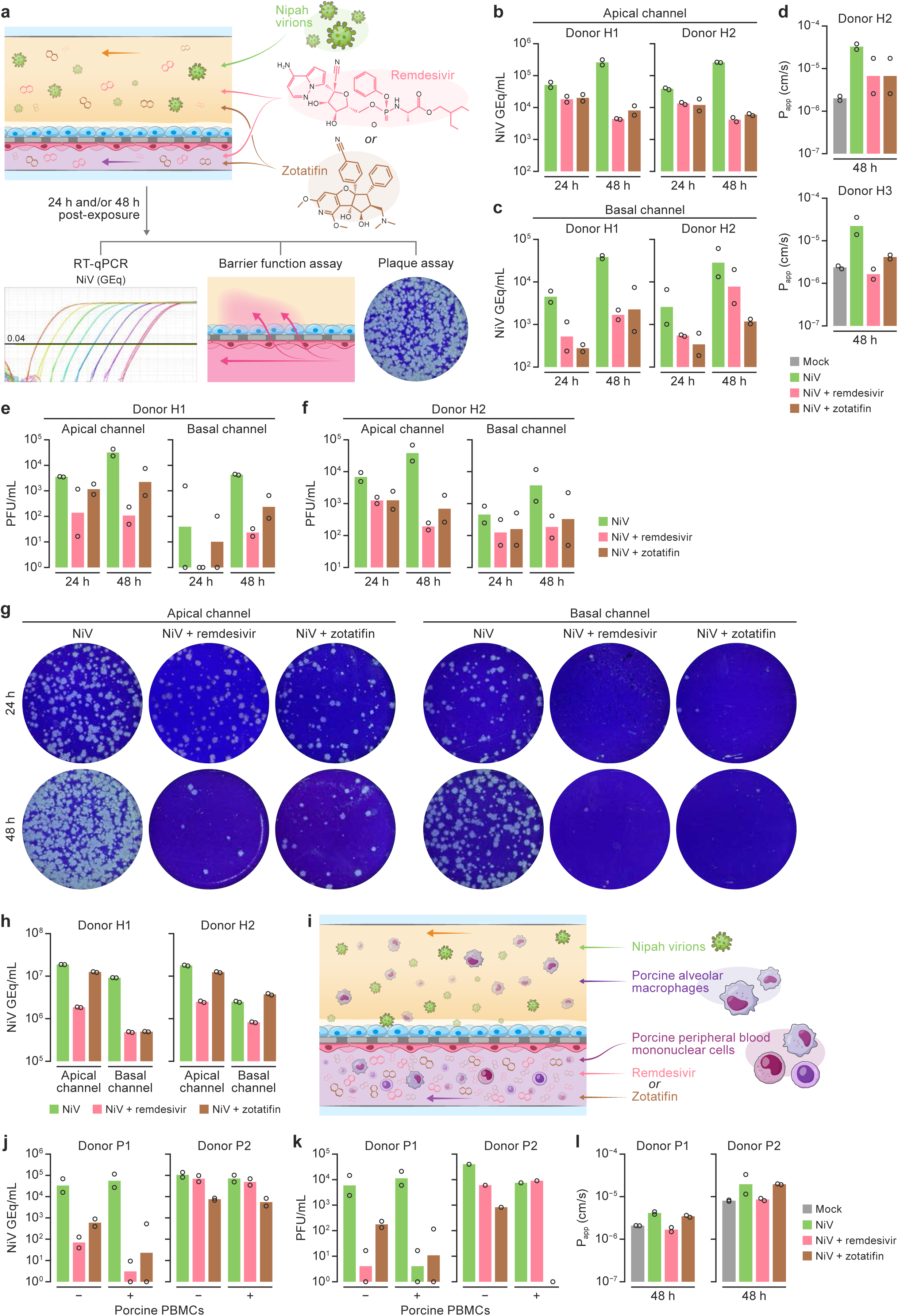
Small-molecule antivirals reduce Nipah virus titers in human lung organ-on-chips. **a**, Schematic of the human small airway organ-on-chip (hsaOOC) experiment: hsaOOCs were prepared in a biosafety level 2 (BSL-2) laboratory using epithelial cells from Donor H1 or Donor H2 (apical channel) and endothelial cells from Donor H3 (basal channel), yielding H1/H3 and H2/H3 hsaOOCs. Once cells were fully differentiated under air–liquid interface (ALI) conditions, organ-on-chips were transferred to maximum (biosafety level 4) containment for exposure to Nipah virus (NiV) or mock-exposed control at 0 h. Duplicate OOCs were used per apical channel donor for all experiments. NiV-exposed hsaOOCs were “treated” with either remdesivir or zotatifin in both the apical and basal channels for 48 h, followed by measuring NiV genome equivalents (GEq) by quantitative reverse transcription polymerase chain reaction (RT-qPCR) (**b–c**), hsaOOC polydimethylsiloxane (PDMS; “interstitial”) membrane barrier function (**d)**, and infectious virus by plaque assay in effluents (**e–f**, representative images in **g**) at 24 and 48 h after treatment initiation. **h,** NiV GEq detected by RT-qPCR in human epithelial cells and human endothelial cells at 48 h post-exposure shown by donor. Data represent means of triplicates and duplicates with standard deviations (SDs) for RT-qPCR and plaque assay, respectively. **i,** Schematic of the porcine alveolus organ-on-chip (paOOC) experiment: paOOCs were prepared in a biosafety level (BSL) 2 laboratory using porcine pneumocytes from Donor P1 or Donor P2 (apical channel) and endothelial cells from Donor P1, yielding P1/P3 and P2/P3 hsaOOCs. Once cells were fully differentiated under air–liquid interface (ALI) conditions, organ-on-chips were transferred to maximum (biosafety level 4) containment and used as described above for hsaOOCs, but with mechanical stretch, antiviral addition only to the basal channels, and addition of porcine alveolar macrophages into the apical channels and porcine peripheral blood monocular cells (PBMCs) into the basal channels. **j,** NiV GEq measured by RT-qPCR. **k,** infectious NiV detected by plaque assay at 48 h in apical channel effluents. **l,** Changes in barrier function shown as permeability measured at 48 h. P_app_, apparent permeability coefficient.

Next, in a similar approach to that used in hsaOOCs, we evaluated antiviral activity in a more complex paOOC system, notable for the shortening of the ALI establishment period to 8 d, administration of antivirals to only the basal channels (thereby mimicking clinical drug injection), and addition of mechanical stretch as well as a cellular immune system component by flowing porcine alveolar macrophages and porcine peripheral blood mononuclear cells through the apical and basal channels, respectively (**Figure 3i**). Similar to the results in hsaOOCs, antiviral “treatment” of paOOCs without the immune cells resulted in a decrease of NiV titers as determined by plaque assay (**Supplementary Figure 1**). Interestingly, this reduction was further enhanced in the presence of the immune cells in P1/P3 paOOCs but not in P2/P3 paOOCs as judged by RT-qPCR results (**Figure 3j**). The plaque assay results for P1/P3 paOOCs were consistent with the RT-qPCR results, but P2/P3 paOOCs plaque assay data indicated NiV reduction with zotatifin but not with remdesivir (**Figure 3k**). By barrier function testing, remdesivir-“treated” paOOCs had reduced barrier permeability compared to zotatifin-“treated” and “untreated” paOOCs (**Figure 3l**).

Together, albeit from a limited number of comparisons, our data indicate that OOCs can be used to evaluate the antiviral activity of small compounds in a more complex setting than standard 2D cell culture.

### Nipah virus exposure of human small airway lung organ-on-chips results in an increase in relevant inflammatory cytokines

NiV exposure of conventionally cultured primary human lung endothelial and epithelial cells induces expression of major inflammatory cytokines, such as C-C motif chemokine ligand 2 (CCL2), C-X-C motif chemokine ligands 8 (CXCL8) and 10 (CXCL10), and interleukin 6 (IL6) (<u>Elvert et al., 2020</u>; <u>Erbar & Maisner, 2010</u>; <u>Lo et al., 2010</u>), which are hypothesized to play important roles in acute lung injury. We therefore evaluated cytokine responses to NiV exposure (as compared to mock exposure) in hsaOOC effluents in the absence or presence of remdesivir or zotatifin. As shown in **Figure 4**, increased levels of these (CCL2, CXCL8, CXCL10, and IL6) and other major inflammatory cytokines, such as interleukin 1A (IL1A), receptor type 1 (IL1R1), were detected in both apical and basal channel effluents of H1/H3 hsaOOCs and H2/H3 hsaOOCs at 24 h after NiV exposure and, with the exception of CCL2, also at 48 h post-exposure. The same cytokines were elevated in the effluents from H2/H3 hsaOOCs, but to different degrees. For instance, IL1R1 levels were largely comparable across all parameters between the two hsaOOCs but IL1A levels were markedly higher at the 24 h time point in basal effluents compared to those of H1/H3 hsaOOCs, and CCL2, CXCL8, and IL6 levels were generally increased. In addition, unlike the effluents of H1/H3 hsaOOCs, levels of epidermal growth factor (EGF) and vascular endothelial growth factor A (VEGFA) were elevated. As expected, we found that antiviral “treatment” impacts these proinflammatory cytokine levels (**Figure 4**). For instance, at 48 h after virus-exposure, “treatment” with both remdesivir and zotatifin significantly reduced IL1A levels in apical channel effluents of both OOCs. On the other hand, reductions in basal channel effluents were only found in H2/H3 hsaOOCs. Remdesivir “treatment” significantly decreased IL1R1 levels in effluents of both hsaOOCs in apical and basal channel effluents at 48 h compared to the NiV-exposed, “untreated” OOCs. CCL2, CXCL8, IL1R1, and IL6 levels decreased significantly in both effluents at both time points after zotatifin “treatment” in both hsaOOCs.

**Figure 4.**
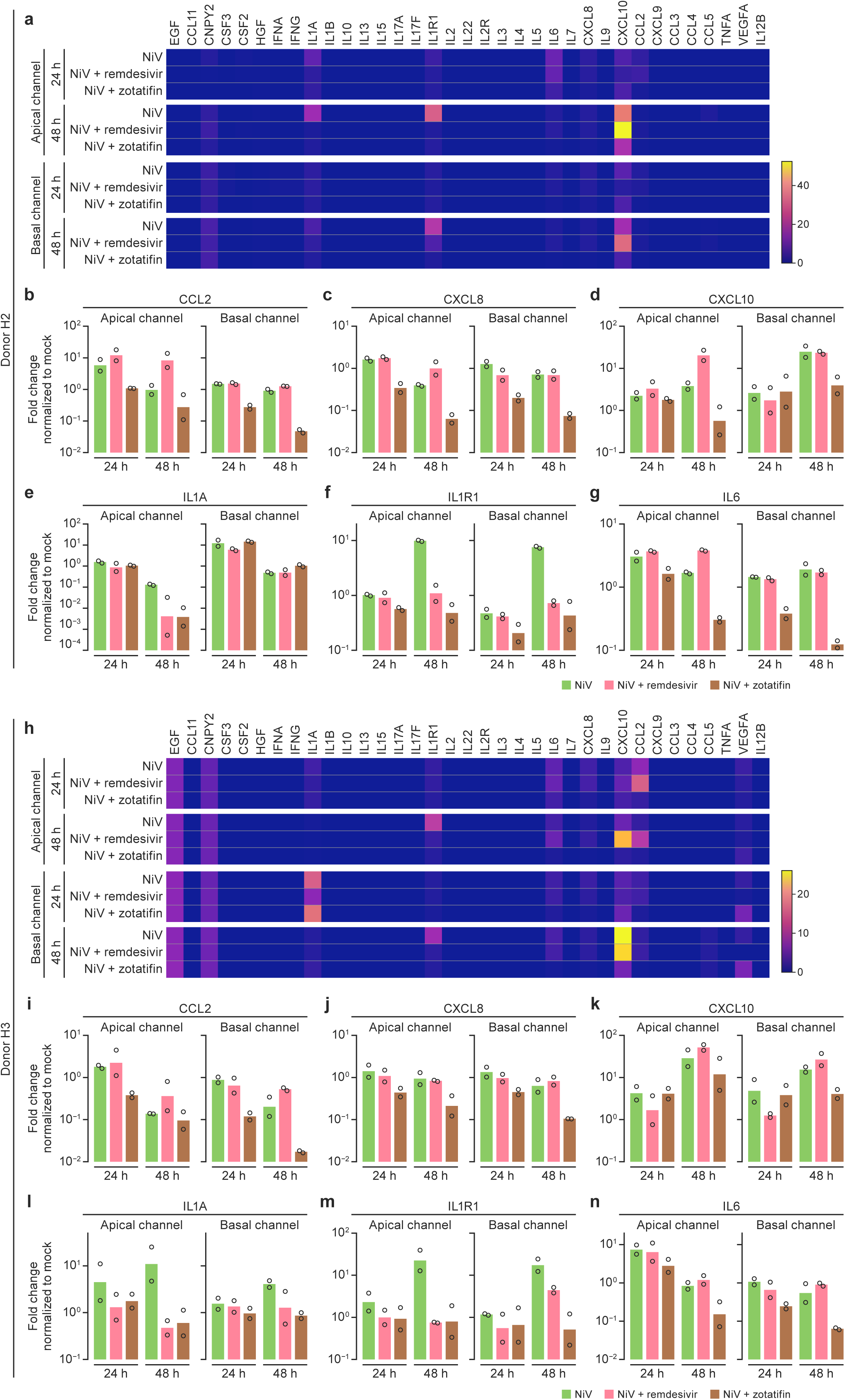
Nipah virus exposure of human small airway lung organ-on-chips results in an increase in relevant inflammatory cytokines. **a and h**,Heat maps showing proinflammatory cytokine levels detected in apical and basal effluents of human small airway lung organ-on-chip (hsaOOCs) seeded with cells from Donor H1 and Donor H3 (H1/H3 OOCs) or Donor H2 and Donor H3 (H2/H3 OOCs) as measured by a 35-plex bioplex assay after NiV exposure in absence of presence of remdesivir or zotatifin. Cytokine names are abbreviated following HUGO Gene Nomenclature Committee recommendations (https://www.genenames.org/). Colors represent relative levels of secreted cytokines from low (blue) to high (yellow). **b–g and i–n,** Selected cytokine quantification in effluents from apical and basal hsaOOC effluents in absence of antiviral “treatment”. Experiments were performed in duplicate.

Together, these data indicate that key proinflammatory cytokine responses to NiV infection in hsaOOCs resemble those in traditional NiV-exposed 2D cultures and *in vivo* models, and that modulation of NiV infection by antivirals produces broadly similar responses that nevertheless vary across cell donors.

### Nipah virus exposure of human small airway lung organ-on-chips induces endothelial activation and subsequent inflammation

In standard tissue culture and transwell systems, NiV infection of microvascular lymphatic lung and brain endothelial cells is associated with transendothelial permeability and cytokine secretion (<u>Erbar & Maisner, 2010</u>; <u>Lo et al., 2010</u>). To evaluate whether these observations are mimicked in NiV infected hsaOOCs, we lysed the human endothelial cells in the basal channels of H1/H3 and H2/H3 hsaOOCs and characterized the gene transcription of 96 human inflammatory markers by RT-qPCR array in the absence or presence of antivirals (**Figure 5a**). As expected, significant changes in gene transcription were measurable in NiV-exposed hsaOOCs compared with nonexposed hsaOOCs in the absence of antivirals. Some genes encoding inflammation markers were upregulated, whereas others were downregulated, but changes varied considerably between H1/H3 and H2/H3 hsaOOCs, suggesting donor-to-donor variability (**Figures 5b–c**). Unsurprisingly, antiviral “treatment” impacted inflammatory gene expression in specific antiviral-dependent and donor-dependent manners (**Figures 5d–e**). Overall, gene expression patterns were generally consistent with cytokine expression (**Figure 4**).

**Figure 5.**
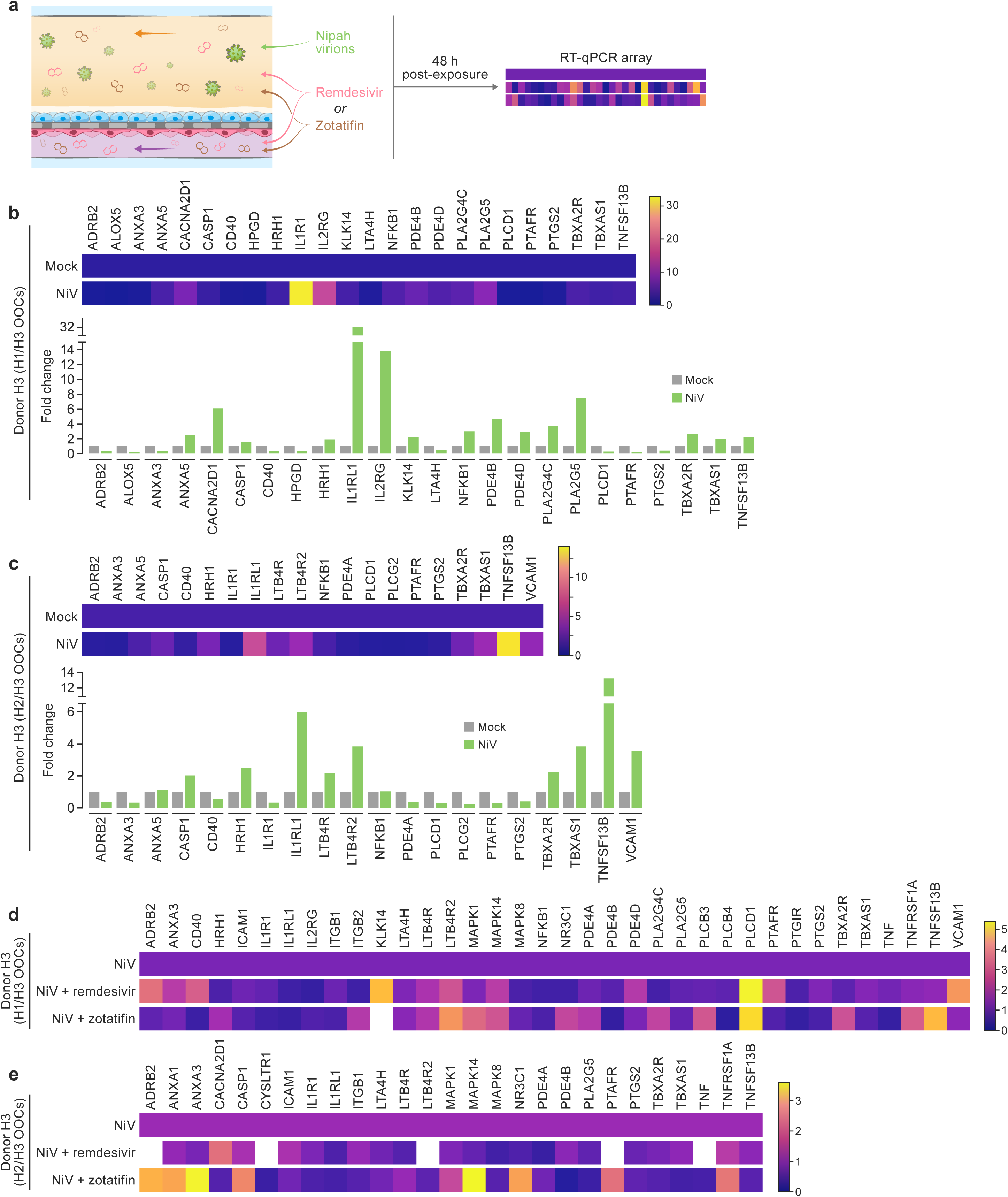
Nipah virus exposure of human small airway lung organ-on-chips induces endothelial activation and subsequent inflammation. **a**, Schematic of the experiment: A quantitative reverse transcription polymerase chain reaction (RT-qPCR) array was used to measure mRNA levels of endothelial cell activation markers induced by Nipah virus (NiV) or mock exposure in the presence and absence of antivirals in human small airway lung organ-on-chips (hsaOOCs). Duplicate hsaOOCs were used per condition. **b and c,** Heat maps and bars showing fold gene induction changes for selected markers in NiV-exposed hsaOOCs compared to mock-exposed hsaOOCs. Colors in the gradient represent relative levels of gene induction from low (blue) to high (yellow). **d and e,** Heat map comparing hsaOOCs exposed to NiV alone compared to hsaOOCs exposed to NiV and “treated” with remdesivir or zotatifin. Fold changes are derived from the mean relative quantification (RQ) values of gene inductions compared to control.

Together, these results demonstrate that the endothelial activation leading to subsequent inflammatory responses to some RG-4 viruses, such as NiV, can be recapitulated in hsaOOCs.

### Nipah-virus-induced neutrophil infiltration can be modelled in human small airway lung organ-on-chips

NiV infection triggers tissue-infiltrative immune responses, including by leukocytes, such as neutrophils, that secrete proinflammatory cytokines, such as CXCL8, CXCL10, and IL6 (<u>Escaffre et al., 2017</u>). As part of our proof-of-principle objective, we isolated (PTPRC^+^+CEACAM1/5/6/8^+^) neutrophils from fresh human blood (**Supplementary Figures 2a–c**) to add this cellular component to the OOC system. We piloted this experimental addition in a BSL-2 laboratory setting: hsaOOCs were exposed to tumor necrosis factor (TNF) for 6 h to mimic infection, then neutrophils were perfused through the hsaOOC basal channel for 2 h. Twenty-four hours later, we detected infiltrating neutrophils in both OOC channels (**Supplementary Figure 2d**). Subsequently, we repeated this experiment in the BSL-4 laboratory, substituting NiV for TNF exposure, and immunostained at 24 h post-exposure for neutrophil marker myeloperoxidase (MPO) and NiV G on the human endothelial and human epithelial cell layers. Neutrophils were detected among cells in both OOC channels, confirming the results from the TNF experiment (**Figures 6a–c**). Cytokine responses in NiV-exposed and nonexposed H1/H3 and H2/H3 hsaOOCs indicated a widespread immune response in the presence of neutrophil flow that was absent in nonexposed or NiV-exposed hsaOOCs without neutrophil flow (**Figures 6d–e**, **Supplementary Figure 3**).

**Figure 6.**
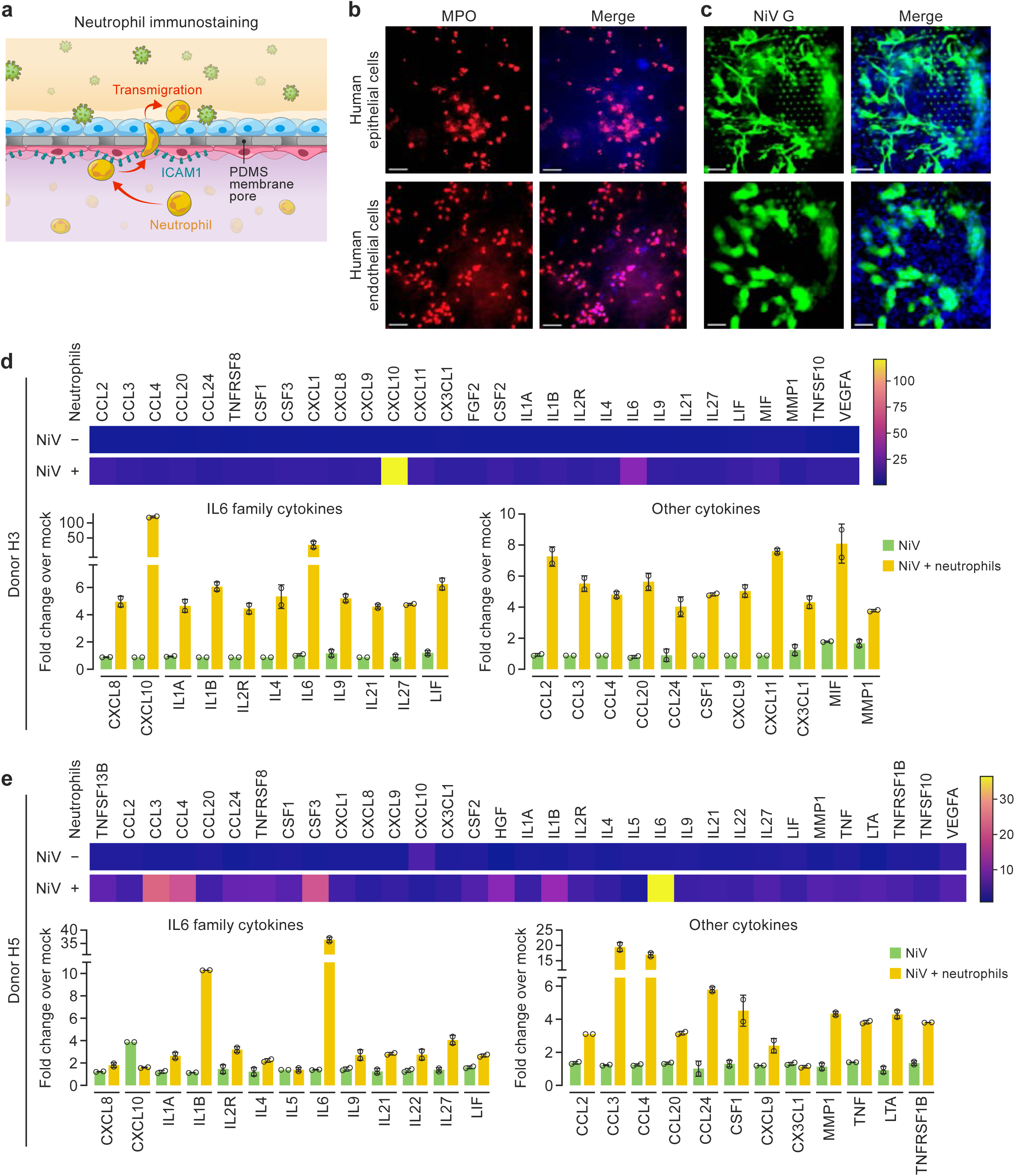
Nipah virus-induced neutrophil infiltration can be modelled in human small airway lung organ-on-chips. **a**, Schematic of the experiment: Neutrophils sorted from fresh human blood were flowed through the basal channels of human small airway lung organ-on- chips (hsaOOCs) after Nipah virus (NiV) exposure. **b,** Neutrophil infiltration, detected by staining neutrophil marker myeloperoxidase (MPO; red), into the apical channels from the basal channels of hsaOOCs 24 h after NiV exposure (*n*=2). **c,** NiV glycoprotein stained with a specific antibody (NiV G; green) (*n*=2). Nuclei (in **b** and **c**) were stained with Hoechst (blue). Scale bar in both panels, 100 µm. **d and e,** Heat maps showing cytokine levels detected in hsaOOC basal channel effluents in NiV-exposed hsaOOCs with or without neutrophils at 24 h post-exposure. Colors in the gradient represent relative levels of secreted cytokines from low (blue) to high (yellow).

Together, these results indicate that neutrophil migration had occurred from the basal into the apical channels of the hsaOOCs across the PDMS (“interstitial”) membrane during NiV infection, suggesting that immune cell migration from the periphery in response to RG-4 virus infection, at least in part, be modeled in OOCs at BSL-4.

## DISCUSSION

Animal experimentation is under increased scrutiny and new U.S. government requests and mandates to complement, reduce, or replace nonhuman animal models (<u>Food and Drug</u> <u>Administration, 2025</u>; <u>National Institutes of Health, 2025a</u>, <u>2025b</u>; <u>U.S. Congress, 2022</u>) compel infectious disease laboratories to include new approach methodologies (NAMs) in their standard research portfolios. Hesitation to do so stems in part from lack of NAM expertise among institute staff, perception that NAMs are premature technologies with little predictive or translational value, and concerns regarding the applicability of NAMs with respect to established safety protocols—particularly in high (biosafety level 3 [BSL-3]) and maximum (BSL-4) containment laboratories.

Here, we demonstrate the operational feasibility and scientific proof of principle that lung OOCs can be used to study a Risk Group 4 (RG-4) host–pathogen interaction in a BSL-4 laboratory. Briefly, we used two types of OOCs emulating the three-dimensional (3D) microenvironments of a human small airway and the porcine alveolus to characterize the effects of Nipah virus (NiV) infection under various conditions. In this work, we strategically addressed several operational challenges, including in our selection of a commercially available OOC system (to reduce the complexity of initial operational standup); the development of novel safety and workflow protocols; and the establishment of mirrored BSL-2 (staff training, assay development, non-NiV experiments) and BSL-4 (NiV experiments) setups (**Figures 1–2**). In the face of prevailing concerns about the considered obstacles to using NAMs in maximum containment settings, we demonstrated that, although challenging, setup is possible and can be implemented by non-NAM experts (albeit with considerable input from experts at the company that made the NAM and colleagues with experience who provided subject matter expertise; see Acknowledgments). Further, we demonstrated that samples taken from OOCs are of sufficient quantity and quality to perform standard virological assays (e.g., cell characterization by immunofluorescence, quantitative reverse transcription polymerase chain reaction [RT-qPCR] and plaque assay), with results mostly in line with previously published results from conventional two-dimensional (2D) *in vitro* and *in vivo* experiments (<u>Elvert et al., 2020</u>; <u>Erbar &</u> <u>Maisner, 2010</u>; <u>Lo et al., 2010</u>) as well as static air–liquid interface (ALI) culture models (<u>Escaffre et al., 2016</u>). For instance, after exposure of human small airway lung OOCs (hsaOOCs) to NiV via the “airway” channel (mimicking respiratory/aerosol exposure), we demonstrated productive NiV infection in primary endothelial and differentiated epithelial cells, indicating NiV migration across the “interstitial” polydimethylsiloxane (PDMS) membrane (**Figure 2**); from these cell layers, we measured cytokine responses generally consistent with those measured previously in 2D monocultures and indicating severe “lung” inflammation (**Figures 4–5**); we reproduced the known anti-NiV activity of remdesivir (<u>Lo et al., 2019</u>) after “inoculation” of the OOC “capillary” channel, mimicking intravenous drug injection (**Figures 3– 5)**; we novelly demonstrated the anti-NiV activity of zotatifin, a small host-targeted molecule that had not been evaluated previously in the context of any NiV infection (<u>Gordon et al., 2020</u>; <u>Müller et al., 2021</u>; <u>Obermann et al., 2022</u>) (**Figures 3–4)**; and, via the addition of neutrophils into the “capillary” channels, we demonstrated that additional immune cellular complexity relevant to the host–virus interaction can be introduced into the lung OOC system (<u>Escaffre et</u> <u>al., 2017</u>) (**Figure 6)**.

Although just a pilot study focused on a single OOC system (and with a limited number of OOCs and donors) and a particular RG-4 infection, our proof-of-principle results encourage further deployment of this emerging technology, including in high-containment settings. Indeed, the potential utility of NAMs to study RG-4 viral infections currently stands at a critical threshold, and community effort will be needed to meet that potential. Within the system described here, challenges fall into three main categories:

1. Choice and source of seed cells: Most OOC efforts rely on acquisition of primary cells from donors via commercial providers. As a result, there will be considerable heterogeneity (and thus intra- and inter-experiment result variation) across institutes and countries, not only regarding origin and genetics of cells, but also due to difference in cell quality. We deliberately used cells from two human and two porcine epithelial cell donors to demonstrate this problem. In addition, we were unable to obtain endothelial cells, epithelial cells, and neutrophils from the same human donor or endothelial cells, epithelial cells, macrophages, and peripheral blood mononuclear cells (PBMCs) from the same porcine donor, thus resulting in “mix-and-match” OOCs;
2. Analysis: The small footprint of our selected OOCs limits the overall number of cells that can be cultured in them. In addition, retrieval of cells from the OOCs is not trivial, and significant cell loss is expected, restricting downstream analyses as compared to *in vivo* experiments that typically generate ample amounts of tissues for collection. In part, we mitigated this issue by also performing analysis *in situ* (i.e., within the OOC) using immunofluorescence staining. However, microscopic assessment of the lung OOCs we used was challenging; for instance, we could not easily distinguish apical and basal channels using the conventional microscopy available in our BSL-4 laboratory. The head piece of the positive pressure suit commonly worn in the BSL-4 laboratory prevents a clear view through a binocular microscope in the absence of a display monitor; even with display access, the considerable time needed for careful review presents a challenge in a BSL-4 setting in which time must be limited and focused on tasks that require the handling of infectious viruses. Ideally, inactivated exposed OOCs would be removed from maximum containment for downstream analyses, including, for example, the imaging of stained OOCs on an Operetta CLS High-Content Analysis System, an instrument that has a large footprint and is thus not suitable for BSL-4 laboratory with limited space. Additionally, the total number of OOCs included in any given experiment was small (up to *n*=20) due to the length and complexity of the protocol requiring collaboration of multiple laboratory staff. Other commercially available OOC systems may offer higher throughput depending on the scientific needs. Future efforts will be enhanced by upgrading hardware in the BSL-4 laboratory, optimizing retrieval methodology, and applying technological advances in “minimal” sample volume or cell number assays (such as microfluidics); and
3. Biosafety: There are currently no across-institute standard operating procedures (SOPs) for “inactivation” of OOCs (i.e., pathogens therein) for downstream analyses outside of high or maximum containment laboratories. Typically, SOPs for *in vitro* work in these laboratories are written specifically for cell monolayers or tissues sampled from animals. OOCs do not fit either of these two categories. In our setting, OOCs were classified as a separate category and, to demonstrate inactivation of cells within the channels, a simple proof-of-principle experiment was performed: Vero E6 cells, an immortalized cell line that is highly permissive to NiV infection, were seeded into both channels of OOCs and exposed to NiV at an multiplicity of infection (MOI) of 1. After 48 h, each channel was flushed with PBS to remove media, and 10% of neutral buffered formalin (NBF) was subsequently pushed through each channel at a volume equal to ten times the volume of what an entire channel holds at a rate of 336 µL/h for 30 min. Subsequently, the OOCs were disconnected from the culture module and dropped into a container filled with 10% NBF. After 10 min, OOCs were removed from containment as approved by instructional biosafety committee. Future efforts in the high containment setting will need careful deliberation by scientists and biosafety experts towards agreement and standardization of inactivation and other biosafety protocols.

Independent of the system described here, research is potentially limited by the types of commercially available OOC systems. We used a virus known to infect the lungs in a relatively well-established lung OOC system; of course, diverse viruses affect numerous organs, many of which cannot yet be modelled in OOC systems. Further complicating matters, several commercial OOC systems are available, and all have platform-specific advantages and disadvantages. Laboratories with small footprints, typical of high and maximum containment facilities, likely cannot simultaneously house multiple systems and thus would either have to select a particular one on which to focus or establish SOPs for frequent system replacement within containment depending on experimental needs.

Many of these challenges could be overcome through inter-institutional collaboration and anticipated near-term scientific breakthroughs. For instance, induced pluripotent stem cell (iPSC) technology can be pivoted to populate future human and nonhuman animal cell banks for standardized creation of well-defined primary seed cells (<u>Koceva & Mosig, 2025</u>). Such an approach would address the heterogeneity and subsequent reproducibility challenges and would also expand opportunities to study the influence of human patient variability (across sexes, ages, ethnicities, and comorbidities) on infectious disease pathogenesis in a manner that cannot be modeled in animals. Sufficiently promising results obtained with OOCs, optimally benchmarked in a species-specific manner, could justify a reduction of animal rooms in containment laboratories making way for more space for OOC operations—and thereby addressing the footprint constraints. Laboratories could work together on SOP optimization, including system, reagent, and assay choices towards independent result replication.

The future for OOCs, including their use in the high and maximum containment settings, looks promising. Even in the system we describe, several immediate improvements are possible, including the addition of mechanical stress (i.e., simulation of breathing via membrane stretching and de-stretching) to the hsaOOCs and possibly the use of true aerosol exposure (rather than simulated with virus-containing medium flow); both additions would move the human platform closer to approximating the *in vivo* lungs in miniature. Small-molecule antivirals (similar to remdesivir and zotatifin) could be delivered through the basal channels using concentrations derived from human or nonhuman pharmacokinetics data, thereby simulating actual drug concentrations over time used *in vivo*. Finally, looking ahead, OOCs simulating different organs could be connected in series (creating effectively a “body-on-a-chip”) to study holistic questions pertaining to pathogenesis or medical countermeasures development.

## ACKNOWLEDGMENTS

We are grateful to Tyler Goralski and Kyle Glover (DEVCOM) for continuous advice and troubleshooting. We are thankful to Elena N. Postnikova (NIAID IRF-Frederick) for providing all compounds, Christopher C. Broder (Microbiology and immunology Department, Uniformed Services University of the Health Sciences, Bethesda, MD, USA) for providing Nipah virus protein G antibody, and Gregory Kocher (NIAID IRF-Frederick) for propagating Nipah virus as required.

This research was supported in part through a Laulima Government Solutions, LLC, prime contract with the U.S. National Institute of Allergy and Infectious Diseases (NIAID) under Contract No. HHSN272201800013C. S.M.B., S.Y., J.G., A.D., S.M.A., B.P., R.B.-C., D.F.P.R., J.W., A.C., and G.W. performed this work as employees of Laulima Government Solutions, LLC. J.P.T. and J.H.K. performed this work as employees of Tunnell Government Services (TGS), a subcontractor of Laulima Government Solutions, LLC, under Contract No. HHSN272201800013C. This work was also supported in part with federal funds from the NIH National Cancer Institute (NCI), under Contract No. 75N91019D00024 with Leidos Biomedical Research, Inc. I.C. performed this work as an employee of Leidos Biomedical Research, Inc., as supported by the Clinical Monitoring Research Program Directorate, Frederick National Laboratory for Cancer Research, sponsored by NCI.

This research was supported in part by the Intramural Research Program of the National Institutes of Health (NIH). The contributions of the NIH author(s) are considered Works of the United States Government. The findings and conclusions presented in this paper are those of the author(s) and do not necessarily reflect the views of the NIH or the U.S. Department of Health and Human Services.

## Author contributions

S.M.B., J.H.K., and G.W. designed the method. S.M.B., J.P.T., S.Y., J.G., A.D., S.M.A., B.P., R.B.-C., D.F.P.R., and B.B. implemented the method and conducted data analysis. S.M.B., S.M.A., B.P., I.C., G.P., N.C.K., J.H.K., and G.W. analyzed the results. S.M.B., J.P.T., S.Y., J.G., J.W., A.C., G.P., N.C.K., J.H.K., and G.W. wrote and edited the manuscript.

## Competing interest

The authors declare no competing interests.

## MATERIAL AND METHODS

### Ethics

For isolation of neutrophils, whole blood samples were drawn at the Integrated Research Facility at Fort Detrick from healthy human volunteers after informed consent and in compliance with U.S. Department of Health and Human Services Code of Federal Regulations (specifically, 45 CFR 46; the “Common Rule”) and institutional regulatory board policy. All other primary cells from human donors were acquired from commercial vendors (**Supplementary Table 1**) and their use did not require institutional review board approval. This work has been reviewed and approved by the National Institutes of Health Institutional Biosafety Committee and Institutional Review Board.

### Cells

Frozen primary human small airway epithelial cells (human epithelial cells) were obtained from two different human donors (Donor H1 and Donor H2), and frozen primary human lung microvascular endothelial cells (human endothelial cells) were obtained from a third human donor (Donor H3; **Supplementary Table 1**). One week prior to seeding cells into human small airway lung organ-on-chips (hsaOOCs), human endothelial cells were cultured and expanded in EGM-2 MV Microvascular Endothelial Cell Growth Medium-2 BulletKit (Lonza; Cat. #CC- 3202) medium in Nunc EasYFlask cell culture flasks (Thermo Fisher Scientific; Cat. #156499). Four days prior to hsaOOC seeding, human epithelial cells were thawed with SAGM Small Airway Epithelial Cell Growth Medium BulletKit (Lonza; Cat. #CC-3118) medium in Nunc EasYFlask cell culture flasks coated with 0.1% Gelatin-Based Coating Solution- ready to use (Cell Biologics; Cat. #CB6950). Gentamicin (GA-1000) in both medium kits was substituted with 1% Penicillin-Streptomycin (Sigma-Aldrich; Cat. #4333-100ML). Media were changed every 2–3 d.

Frozen primary porcine alveolar epithelial cells (porcine pneumocytes) were obtained from two different cell donors (Donor P1 and Donor P2), and frozen primary porcine lung microvascular endothelial cells (porcine endothelial cells) were obtained from a third donor (Donor P3) (**Supplementary Table 2**). For seeding into porcine alveolus lung organ-on-chips (paOOCs), porcine cells were thawed and expanded in the same fashion as the human primary cells using Complete Endothelial Cell Medium /W Kit (Cell Biologics; Cat. #M1168) medium and Complete Epithelial Cell Medium /W Kit (Cell Biologics; Cat. #M6621) medium for porcine endothelial cells and porcine pneumocytes, respectively.

### Lung organ-on-chips cell seeding/differentiation and establishment of air–liquid interfaces

hsaOOCs were set up in accordance with vendor instructions, with some simplifying modifications. Briefly, microfluidic chips (Fisher Scientific; Cat. #NC2084454) were activated using kit-provided ER1 and ER2 solutions (Fisher Scientific; Cat. #NC1624644 and NC2082596, respectively) under ultraviolet light for 15 min. The apical (“parenchymal”/“airway”/top) channels were coated with Human Collagen Type IV (Millipore Sigma; Cat. #C5533) and the basal (“vascular”/“capillary”/bottom) channels were coated with Human Collagen Type IV and Corning Fibronectin (Corning; Cat. #356008) for up to 3 d and stored at 4°C. One hour prior to cell seeding, the activated chips were warmed by storing at 37°C for at least 1 h. Donor H3 human endothelial cells were seeded into the basal (“vascular”/“capillary”) channel at a density of 3–5 × 10^6^ cells per mL. The hsaOOCs were then inverted and incubated for 2–4 h at 37°C to facilitate cell attachment. Following confirmation of cell attachment by brightfield microscopy, the hsaOOCs were then reverted to face upward, and the apical (“parenchymal”/“airway”/top) channels were seeded with the human epithelial cells from either Donor H1 or Donor H2 at a density of 3 × 10^6^ cells per mL (resulting in H1/H3 or H2/H3 hsaOOCs). After 20–24 h, the hsaOOCs were connected to the pod reservoirs, which served as media reservoirs, and then inserted into a Zoë-CM2 Culture Module (Emulate). Via the culture module, the flow rate was set to 45 µL/h rate for both channels and the respective media flowed through the channels for 48 h until cell layers were judged confluent using bright field microscopy. To set up the air–liquid interface (ALI), apical channel media were replaced under the same flow rate with PneumaCult-ALI Medium (STEMCELL Technologies; Cat. #05001) supplemented with Hydrocortisone Stock Solution (STEMCELL Technologies; Cat. #07926) at a concentration of 0.48 µg/mL, Heparin Solution (STEMCELL Technologies; Cat. #07980), and Epidermal Growth Factor (EGF), Human Recombinant (ProSpec Protein Specialists; Cat. #CYT- 332) (from here on: complete ALI medium). After 5–6 d, ALI conditions were established to induce human epithelial cell differentiation by removing all media from apical channels and subsequently setting the flow rate to zero (“air exposure”) while flowing a 1:1 mixture of complete ALI medium and EGM-2 MV Microvascular Endothelial Cell Growth Medium-2, supplemented with the synthetic retinoid EC 23 (Tocris Bioscience; Cat. #4011) to a concentration of 5 µM (from here on: complete human endothelial medium) through the basal channels at 45 µL/h. Mucus was removed twice per week by flushing Corning Dulbecco’s Phosphate-Buffered Saline, 1X without calcium and magnesium (DPBS; Corning; Cat. #21-031- CV) through the apical channels at 1,000 μL/h for 3 min, incubating the epithelial cells with DPBS for 10 min in static (no-flow) conditions, and then clearing the apical channels of DPBS to re-establish the ALI conditions. The basal channel media were replenished three times per week.

paOOCs were set up similarly to hsaOOCs with minor modifications: The apical and basal channels were both coated with Human Collagen Type IV and Laminin from human placenta (Sigma-Aldrich; Cat. #L6274). Porcine endothelial cells were seeded at a density of 8 × 10^6^ cells per mL and porcine pneumocytes were seeded at a density of 1.6 × 10^6^ cells per mL. To enhance barrier function, Complete Epithelial Cell Medium /W Kit medium was supplemented with 1 μM dexamethasone 3 d after cell seeding. Two days after ALI was established, fetal bovine serum (FBS) was reduced in the basal channel media to 0.5%. Two days after reduction of FBS in media, mechanical stretch was initiated by setting stretch to 5% and frequency to 0.25 Hz and was applied for 4 d before OOCs were judged ready for antiviral “treatment”.

### Immunofluorescence assays prior to organ-on-chip seeding

Cells in 96-well plates were fixed with 100 μL of Formalin, 10% neutral buffered formalin (NBF; StatLab; Cat. #28600-1) for 30 min at ambient temperature and washed with 100 μL of DPBS three times. Each well then received 100 μL of permeabilizing solution, comprised of 0.1% Triton X-100 (Millipore Sigma; Cat. #X100-100ML) in DPBS, followed by an incubation of 5 min at ambient temperature. Cells were washed with DPBS three times and 100 μL of blocking buffer, composed of 1% Bovine Serum Albumin (BSA; Sigma-Aldrich; Cat. #A1933-25G), 10% Donkey Serum (Millipore Sigma; Cat. #D9663) and 0.1% Triton X-100 in DPBS, were added to each well and incubated for 1 h at ambient temperature. Blocking buffer was removed and 100 μL of primary antibodies (**Supplementary Table 3**) diluted in blocking buffer were added for overnight incubation at 4°C. Cells were then washed three times with DPBS, and 100 μL of secondary antibodies (**Supplementary Table 3**), prepared in 1% BSA in DPBS, were added to the cells and incubated for 1 h at ambient temperature. Cells were again washed three times with DPBS and then counterstained with 100 μL of Hoechst 33342, trihydrochloride trihydrate (Hoechst; Invitrogen; Cat. #H3570) diluted in DPBS for 5 min. Counterstain was removed and cells were washed three times in DPBS. Cells were stored in DPBS until high-content imaging was performed using an Operetta CLS High-Content Analysis System with Harmony software (PerkinElmer).

### Virus exposures

#### Stock virus

Nipah virus (NiV; *Paramyxoviridae*: *Henipavirus nipahense*), isolate Bangladesh 2004 (GenBank Accession #AY988601), a Risk Group 4 (RG-4) agent, (<u>U.S. Department of Health</u> <u>and Human Services et al., 2020</u>) was obtained from the U.S. Centers of Disease Control and Prevention (CDC). NiV was propagated in a biosafety level 4 (BSL-4) laboratory by inoculating grivet (*Chlorocebus aethiops* (Linnaeus, 1758)) kidney epithelial Vero E6 cells (BEI Resources; Cat. #NR596), maintained in Dulbecco’s Modified Eagle Medium (DMEM; Gibco; Cat. #11995040) containing 2% heat-inactivated FBS (Millipore Sigma; Cat. #F4135) at a multiplicity of infection (MOI) of 0.01. Cell-culture supernatant was collected 4 d after virus exposure and processed according to institutionally approved standard operating procedures (SOPs) for virus collection. A plaque assay was performed to determine the titer of the stock that was generated following a published protocol (<u>Jensen et al., 2018</u>).

#### Organ-on-chip virus exposures

All virus exposure experiments were performed in a BSL-4 laboratory. Prior to virus exposure, the OOC basal channel inlet reservoirs were replenished with fresh complete human endothelial medium (for hsaOOCs) or Complete Endothelial Cell medium (for paOOCs). A volume of 0.5 mL of NiV diluted in complete ALI medium (for hsaOOCs) or Complete Epithelial Cell Medium /W Kit medium (for paOOCs), calculated to achieve an MOI of 1, were then added to the apical channel inlet reservoirs. With the apical channel flow rate set at 400 μL/h and the basal channel flow rate set at 0 μL/h, NiV was then flowed through the apical channels for 30 min. Then, flow was stopped for static NiV exposure for an additional 1.5–2 h, totaling to 2–2.5 h of virus exposure. Media were then removed from both channels by extracting media from all pod reservoirs and flushing both channels at 1,000 μL/h for 2 min, followed by aspirating any residue media from the apical and basal outlets. The cells were washed by adding 1 mL of DPBS to the inlet and 0.5 mL of DPBS to the outlet reservoirs of the apical channels and flowing it through at 1,000 μL/h for 3 min. DPBS was removed from the reservoirs and the apical channels were cleared of any residue DPBS by flushing them at 1,000 μL/h for 2 min. For hsaOOCs, complete ALI medium was reintroduced to the apical channels, and the basal channels were replenished with complete human endothelial medium. Then, flow was restored at 45 μL/h for both channels. For paOOCs, Complete Endothelial Cell medium (0.5% FBS) was reintroduced to the basal channels, but the apical channel cells were exposed to air (flow rate: 0 μL/h), thereby re-establishing ALI. The flow rate was to 45 μL/h for the basal channels and mechanical stretch was applied at 5% and 0.25 Hz. The diluted virus inocula were recovered from the apical channels as a positive control for downstream assays.

#### Antiviral “treatment”

Remdesivir was provided by OyaGen. Zotatifin (eFT226; Cat. #HY-112163) was purchased from MedChem Express. For antiviral “treatment”, OOCs were exposed to NiV as described above with antivirals present in the basal channel replenishing media (hsaOOCs and paOOCs) and in the NiV exposure medium (hsaOOCs only). Antivirals were again added to the respective media after NiV inocula removal and continuously flowed through the apical and basal channels (hsaOOC) or just the basal channels (paOOC).

### Post-exposure assays

#### Quantitative reverse transcription polymerase chain reaction (RT-qPCR) assay

Effluents from both OOC channels were collected in the BSL-4 laboratory at 24 and 48 h after NiV exposure and stored at -80°C until use. Cells were lysed in their channels at 48 h after NiV exposure. Effluents or cells obtained from OOCs were mixed with TRIzol LS reagent (Thermo Fisher Scientific) per the manufacturer’s instructions to inactivate NiV. RNA extraction from effluents and lysed cells was performed using a MagMAX Viral Pathogen Nucleic Acid Isolation Kit (Thermo Fisher Scientific; Cat. #A48310) with a Kingfisher instrument (Thermo Fisher Scientific) following the manufacturer’s instructions (<u>Bhosle et al., 2018</u>). The forward primer (5’ GTACTCAACCATGAATGAACAGTTG 3’) was designed to recognize nucleotides 8,658– 8,682 in the 5’ untranslated region of NiV’s fusion (*F*) gene, and the reverse primer (5’CTTTAAAGGACACAGTTTAATATCCAATG3’) was designed to recognize nucleotides 8,735–8,763 in the 3’ region of NiV’s glycoprotein (*G*) gene, thereby amplifying the *F–G* intergenic region. The hydrolysis probe, with a 6-carboxyfluorescein (FAM) reporter with a double quencher (FAM-5’CTTAGGACCCAGGTCCATAA3’), was designed to span nucleotides 8,713–8,732 in the 3’ region of the NiV *G* gene.

For RT-qPCR assays, a reaction mixture was prepared with 500 nM of primers, 250 nM of the probe, 1X TaqPath 1-Step RT-qPCR Master Mix, CG (Thermo Fisher Scientific; Cat. #A15299) and Nuclease Free Water (Integrated DNA Technologies; Cat. #11-04-02-01). For each reaction, 5 μL of the extracted RNA were added to 15 μL of the reaction mixture. The completed mixtures were subjected to complementary DNA (cDNA) synthesis (2 min at 25°C, followed by 15 min at 50°C), initial denaturation (2 min at 95°C), 40 cycles of subsequent denaturation (every 10 s at 95°C), annealing/elongation (30 s at 60°C), and final elongation (2 min at 72°C) in a QuantStudio 7 Flex Real-Time PCR System, 96-well, laptop (Thermo Fisher Scientific; Cat. #4485688). RNase-free water (Integrated DNA Technologies; Cat. #11-04-02- 01) was used as the non-template (mock) control. RT-qPCR results were analyzed on QuantStudio 7 software with a threshold of 0.04.

#### Plaque assay

Effluents from both OOC channels were collected in the BSL-4 laboratory at 24 and 48 h after NiV exposure and stored at -80°C until use. Plaque assays were performed with dilutions of each effluent with a slight modification to a previously published protocol (<u>Cong et al., 2017</u>). Vero E6 cells were seeded in 6-well plates 1 d before the assay. Gibco low-glucose DMEM with GlutaMAX (Thermo Fisher Scientific; Cat. #10567014) was used for dilution from 10^-1^–10^-6^. Cells were incubated for 60±10 min at 37°C after added samples, rocking gently approximately every 10–15 min. FMC BioPolymer as DBA Novamatrix Avicel RC-591 overlay (Fisher Scientific; Cat. #NC0821602) and 2X plaque assay medium (Gibco; Cat. #11935046) at a ratio of 1:1 in a total volume of 2 mL were added to each well. Plates were swirled gently, covered, and then incubated for 72 h at 37°C. After incubation, plates were swirled to loosen the overlays, and the overlays were removed carefully. Then, 1–2 mL of 10% NBF containing 0.2% Gentian Violet, 2% (w/v) (RICCA; Cat. #32334) were added to each well. Plates were incubated for at least 30 min at ambient temperature, followed by gentle rocking. Subsequently, the dye was aspirated, and the plates were rinsed thoroughly with tap water and dried at ambient temperature. Then, plaques were counted manually on a lightbox. Viral titers were calculated as PFU per mL.

#### Immunofluorescence assays prior to organ-on-chip seeding

Cells in OOCs were fixed via an Institutional Biosafety Committee-approved protocol by flowing ten times the total channel volume (10 × 33.6 = 336 μL) of 10% NBF through both channels for 30 min at a flow rate of 336 μL/h. The OOCs were then disconnected from the pod reservoirs and fully submerged in 10% NBF to decontaminate the surface of the OOCs for an additional 10 min. OOCs were removed from the fixative and both channels were flushed by pipetting 200 µL of DPBS through both channels and then fully submerged and stored in DPBS. For immunofluorescence assays, cells in both channels were incubated with 100 µL of permeabilizing solution at ambient temperature for 5 min. After incubation, cells were washed three times with 200 µL of wash buffer, which was comprised of 1% Polysorbate 20 (Fisher Scientific; Cat. #BP337-500) in DPBS. The cells were then blocked with 100 µL of blocking buffer for 1 h at ambient temperature. After removing the blocking buffer, cells were incubated with 100 µL of primary antibodies (**Supplementary Table 3**) prepared in blocking buffer overnight at 4°C. Cells were then washed as described previously and incubated with 100 µL of secondary antibodies (**Supplementary Table 3**) prepared in 1% BSA in DPBS for 1 h at ambient temperature. Cells were then washed and counterstained with 1:1,000 diluted Hoechst or with SYTOX Deep Red Nucleic Acid Stain, for fixed/dead cells (Thermo Fisher Scientific; Cat. #S11380) for nuclei visualization for 5–10 min. Finally, cells were washed and the OOCs were stored in ProLong Diamond Antifade Mountant (Invitrogen; Cat. #P36961) until high-content imaging was performed using an Operetta CLS High-Content Analysis System with Harmony software.

#### Barrier function test

OOC barrier permeability was measured with a barrier function assay at 48 h after NiV exposure with a slight modification to a previously published protocol (<u>Si et al., 2021</u>). Briefly, 10 mg Cascade Blue 3kDa fluorescent dye (Invitrogen; Cat. #D7132) were resuspended in 1 mL of sterile cell-culture grade water (Corning; Cat. #5-055-CI) to a working concentration of 10 mg/mL. The tracer medium was prepared by diluting the Cascade Blue 3kDa in complete human endothelial medium (for hsaOOCs) or Complete Endothelial medium (for paOOCs) to final concentrations of 100 µg/mL. After media were removed from the inlet reservoirs, cells in both OOC channels were washed by adding 1 mL of DPBS each into the apical and basal inlet reservoirs and flowing it through the channels at a rate of 1,000 μL/h for 3 min. DPBS was then cleared from all reservoirs and the OOCs were subjected to another 1,000 μL/h flow for 3 min to clear both channels of DPBS. Afterwards, 1 mL of complete ALI medium (for hsaOOCs) or 1 mL of Complete Epithelial medium (for paOOCs) were added to the apical channel inlet reservoirs and 1 mL of the tracer medium was added to the basal channel inlet reservoirs. The flow rate was then set to 600 µL/h for 5 min to prime both channels of the proper media and then reduced to 120 µL/h for 2.5 h. The effluents from both channels were collected and stored at - 80°C until further use. The collection from each channel were aliquoted in triplicates into Cell Culture Microplate, 96 Well, PS, F-Bottom plates (Greiner Bio-One; Cat. #655946) and spectral absorbance (excitation 400 nm/emission 420 nm) was measured on a Spark 20M Multimode Reader (Tecan). The apparent permeability coefficient (P_app_ [cm/s]) was calculated using the formula P_app_ = J / (A × ΔC), in which J (mg/s) is the net dye flux across the membrane/the rate of permeation, A (cm^2^) is the surface area of the membrane through which the dye is diffusing, and ΔC (also sometimes called C_0_) is the initial mass (mg) of the dye in the sampled compartment, (i.e., the basal channel).

#### Proinflammatory cytokine quantification

Effluents from OOC channels were analyzed using a Cytokine 35-Plex Human Panel (Thermo Fisher Scientific; Cat. #LHC6005M) or ProcartaPlex Human Immune Monitoring Panel, 65-Plex (Invitrogen; Cat. #EPX650-10065-901) bioplex assay in duplicates in accordance with the manufacturers’ instructions. Briefly, kit-provided 1X antibody beads were added to 96-well plates, which were then washed twice with 1X washing solution. Kit-provided standards or samples (50 µL) and incubation buffer (25 µL) were added to the wells. Then, the plates were sealed, wrapped with foil, and incubated for 16–18 h at 4°C (35-plex assay) or 2 h at ambient temperature (65-plex assay) on a plate shaker for gentle agitation. The next day, biotinylated detector antibody was added to each well, and the plates were incubated for 30–60 min with gentle agitation at ambient temperature. The plates were washed, and 50 µL of Streptavidin R- PE solution (Thermo Fisher Scientific; Cat. #SA10041) were added to each well. The plates were incubated for 30 min at ambient temperature with gentle agitation and then washed three times with kit-provided wash buffer. Next, 120–150 µL of wash solution were added, and the plates were incubated for 5 min at ambient temperature on a plate shaker for gentle agitation. The plates were read on a Luminex FLEXMAP 3D with xPONENT 4.2 (R&D Systems; Cat. #FLEXMAP-3D-RUO), and analyte concentrations were determined using the Bio-Plex Results Generator 3.0 application in Bio-Plex Manager 6.2 Standard Software (Bio-Rad) by plotting the standard curves for each analyte.

#### RT-qPCR array

An RT-qPCR array for detection of human inflammation marker mRNA transcripts was performed with a TaqMan Array Human Inflammation Panel (Thermo Fisher Scientific; Cat. #4378707) and a 384-well microfluidic card in accordance with the manufacturer’s protocol. cDNA was prepared from the OOC cells after RNA extraction as described above with SuperScript IV VILO Master Mix (Thermo Fisher Scientific; Cat. #11766050). Each reaction contained 16 µL of RNA template and 4 µL of SuperScript IV VILO Master Mix, and cDNA was synthesized by primer annealing (25°C for 10 min), RNA reverse transcription (50°C for 10 min), and enzyme inactivation (85°C for 5 min), followed by a holding step at 4°C. cDNA was quantified using a NanoDrop 8000 Spectrophotometer (Thermo Fisher Scientific; Cat. #1216). For each sample, approximately 25 ng of cDNA were mixed with 55 µL of TaqMan Fast Advanced Master Mix for qPCR (Thermo Fisher Scientific; Cat. #4444557). The prepared samples were loaded onto the array card and centrifuged according to the manufacturer’s instructions to distribute the samples into the wells. The assay was run using the QuantStudio 7 Flex Real-Time PCR System, 96-well, laptop under the following conditions: uracil-DNA glycosylases incubation (50°C for 2 min), enzyme activation (92°C for 10 min), and 40 cycles of denature and anneal/extend (60°C for 20 s and 95°C at 1 s, respectively). RT-qPCR array results were analyzed on design and analysis software (Thermo Fisher Scientific) by removing the outliers and normalizing the data with four endogenous cell markers (18S, ACBT, B2M, and GAPDH) to perform relative quantification with relative thresholds settings.

#### Neutrophil experimentation

Fresh whole blood samples (8,000–25,000 µL) were drawn in compliance with 45 CFR 46 (the “Common Rule”) and institutional regulatory board policy. To isolate neutrophils, blood was treated with ACK Lysing Buffer (Quality Biological; Cat. #118-156-101) for 10 min at ambient temperature in the dark to lyse erythrocytes, centrifuged at 550 x*g* for 5 min, decanted, and gently rocked to resuspend the remaining cells, followed by washing with MACSQuant Tyto Running Buffer (Miltenyi Biotec; Cat. #130-107-207). Samples were blocked with 30 µL per mL of whole blood of Human TruStain FcX (Fc Receptor Blocking Solution) (BioLegend; Cat. #422302) and incubated in the dark for 10 min on ice. Next, a surface-stain master mix (30 µL of PTPRC: APC or FITC, 30 µL of CEACAM1/5/6/8: FITC or APC, 12 µL of Live/Dead Fixable Aqua per mL of whole blood) was diluted in PBS, added to all sample tubes, and incubated in the dark for 20–30 min on ice. Following incubation, samples were washed with MACSQuant Tyto Running Buffer, centrifuged at 550 x*g* for 5 min, decanted, and gently rocked to resuspend cells. Cells were counted and resuspended in MACSQuant Tyto Running Buffer to a concentration of 2.0–4.0 × 10^6^ live cells per mL, passed through 20 µM Pre-Separation Filters (Miltenyi Biotec; Cat. #130-101-812) and loaded into MACSQuant Tyto Cartridges (Miltenyi Biotec; Cat. #130-104-791) for cell sorting with a MACSQuant Tyto Cell Sorter (Miltenyi Biotec; Cat. #130-103-931). A sample was taken at each of the following stages of the process using flow-cytometry acquisition with a 5-laser Cytek Aurora spectral flow cytometer (Cytek Biosciences): pre-stained, pre-sorted, positive-sorting fraction, and negative-sorting fraction. Spectral unmixing was performed in SpectroFlo software (Cytek Biosciences) using unstained cell samples, and single antibodies were incubated with UltraComp eBeads Spectral Unmixing Beads (Thermo Fisher Scientific; Cat. #U20250) and ArC Amine Reactive Compensation Beads (Molecular Probes; Cat. #A1628). Each sort resulted in a 90% or greater population of PTPRC^+^+CEACAM1/5/6/8^+^ cells, with a viability of ≥98%.

A pilot experiment was performed in a BSL-2 laboratory and involved exposing hsaOOCs to 50 ng/mL of Recombinant Human TNF-alpha Protein (TNF; R&D; Cat. #210-TA) for 6 h. Isolated neutrophils resuspended in complete human endothelial medium at a density of 5 × 10^6^ cells per mL were then flowed through the hsaOOC basal channels for 2 h at a rate of 45 µL/h. Neutrophil-containing media were then replaced with regular complete human endothelial medium, and cells were fixed 24 h later as described above. This experiment was repeated in a BSL-4 laboratory using the same conditions with NiV exposure as described above in lieu of TNF. hsaOOCs were either fixed for immunofluorescence assay analysis as described above 24 h after NiV exposure or effluents were collected at 24 and 48 h for RT-qPCR and proinflammatory cytokine quantification assays.

#### Statistical analysis

Data are presented as means of duplicate or triplicate values with standard deviation. All analyses were performed on Prism Version 9.3.1 software (GraphPad).

## Data availability

All data used in this study are in the public domain and available on request.

## Code availability

No code was written for this work.

## Supplementary Figures

**Supplementary Figure 1.**
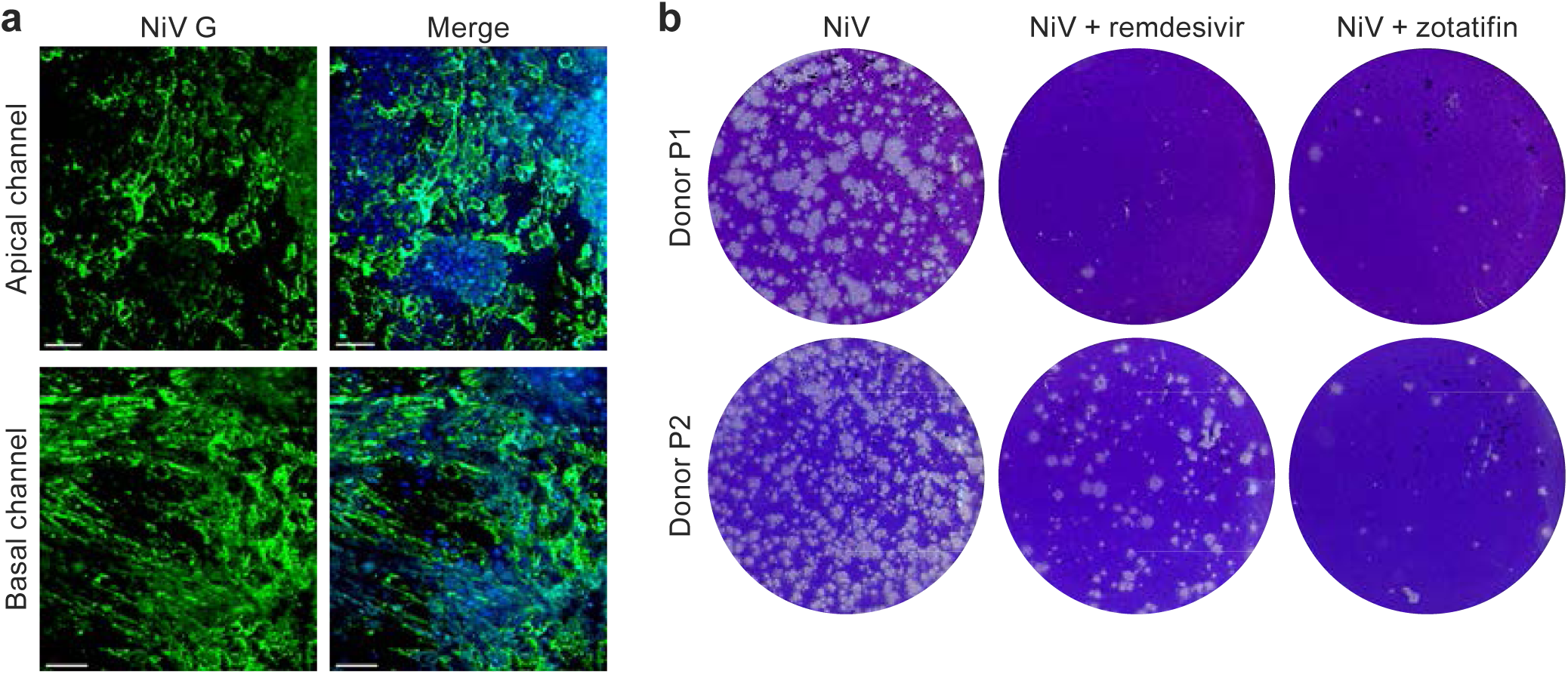
Small-molecule antivirals reduce Nipah virus titers in lung organ-on-chips: Porcine alveolus lung organ-on-chips. **a**, Immunostaining of porcine pneumocytes (apical channel) and endothelial cells (basal channel) for Nipah virus (NiV) glycoprotein (G; green) and nuclei with Hoechst (blue). **b,** Representative NiV plaque assay enumerating virions in effluents collected from “untreated” and antiviral-“treated” porcine alveolus lung organ-on-chips.

**Supplementary Figure 2.**
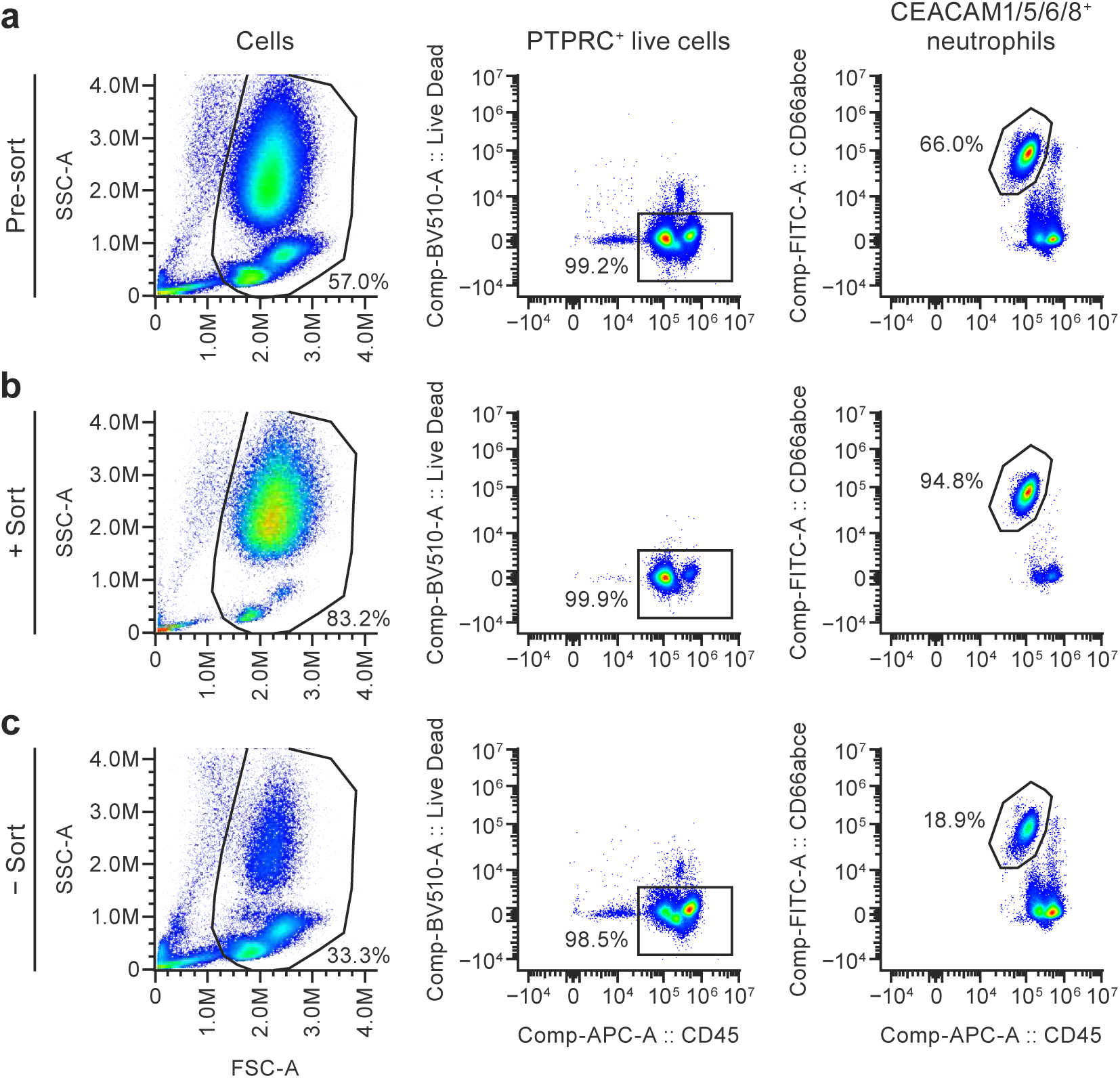
Nipah virus-induced neutrophil infiltration can be modelled in human small airway lung organ-on-chips. Freshly obtained human blood cells were stained to detect leukocyte common antigen, i.e., protein tyrosine phosphatase, receptor type, C (PTPRC) and CEA cell adhesion molecules 1/5/6/8 (CEACAM1/5/6/8) and sorted. The presorted (Pre-sort) (**a**), positively sorted (+ Sort) (**b**), and negatively sorted (- Sort) (**c**) cells were analyzed to detect neutrophil population percentages. The side-scatter cell population was gated and percentages of PTPRC^+^+CEACAM1/5/6/8^+^ neutrophils are shown in presorted, positively sorted, or negatively sorted groups.

**Supplementary Figure 3.**
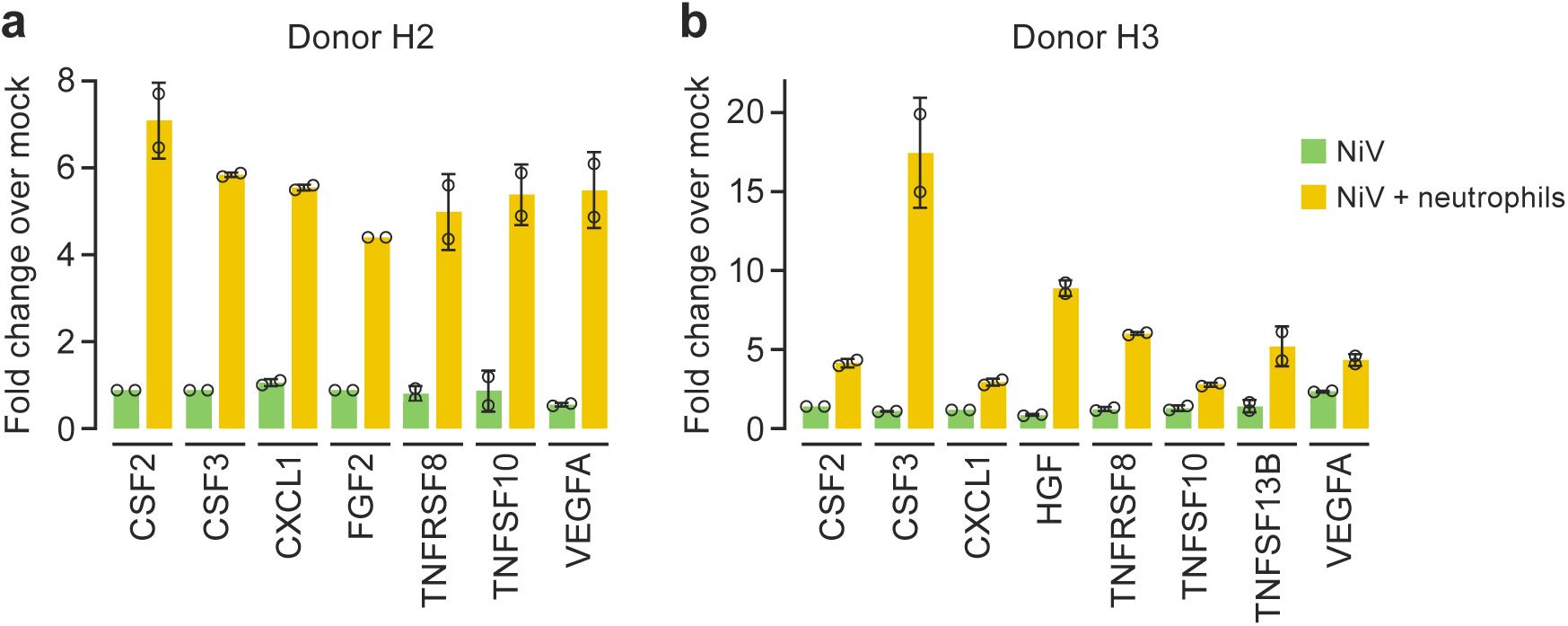
Nipah virus-induced neutrophil infiltration can be modelled in human small airway lung organ-on-chips: immune response in the presence of infiltrating neutrophils. Cytokine concentrations in basal small airway lung organ-on-chip (hsaOOC) effluents at 24 h after Nipah virus (NiV) exposure in absence or presence of neutrophils. CSF2, colony stimulating factor 2; CSF3, colony stimulating factor 3; CXCL1, C-X-C motif chemokine ligand 1; FGF2, fibroblast growth factor 2; HGF, hepatocyte growth factor; TNF, tumor necrosis factor; TNFRSF8, TNF receptor superfamily member 8; TNFSF10, TNF superfamily member 10; TNFSF13B, TNF superfamily member 13b; VEGFA, vascular endothelial growth factor A.

**Supplementary Figure 4.**
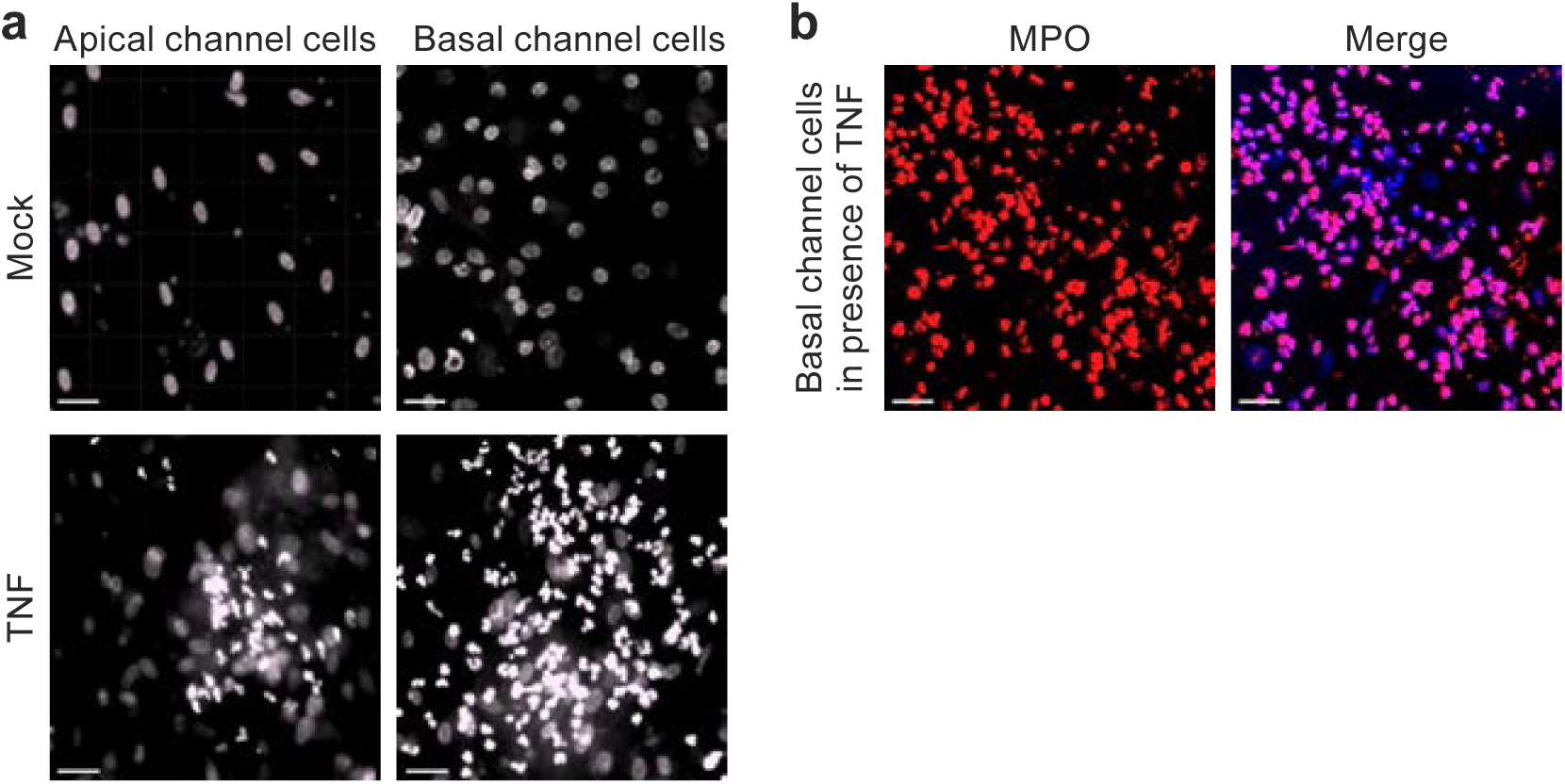
Neutrophil infiltration in cells from apical and basal channels of human small airway lung organ-on-chips after tumor necrosis factor alpha (TNFA) treatment. (a) Small airway lung organ-on-chip (hsaOOC) treated with TNFA showed infiltrating neutrophils in cells from the apical and basal channels at 24 h. Bright field images were captured at 20X with EVOS7000 microscope; scale bar, 40 µm. b, Cells from basal channel stained for MPO1 (red). A merged image with nuclei (blue) is shown; scale bar, 40 µm. MPO, myeloperoxidase; TNF, tumor necrosis factor.

## Supplementary Tables

**Supplementary Table 1.** Metadata for human donors of primary small airway epithelial and primary lung microvascular endothelial cells.

| Human donor | Cell type | Lot number | Sex | Ethnicity | Age (yr) | Weight (kg) | Diabetes | Heart disease | Hypertension | Alcohol use | Smoking |
| --- | --- | --- | --- | --- | --- | --- | --- | --- | --- | --- | --- |
| H1 | Human Small Airway Epithelial Cells (Lonza; Cat. #CC-2547) | 21TL019234 | Female | Caucasian | 78 | 177 | No | Unknown | Yes | No | No |
| H2 | Human Small Airway Epithelial Cells (Lonza; Cat. #CC-2547) | 22TL018267 | Male | Caucasian | 21 | 210 | No | No | No | No | No |
| H3 | HMVEC-L – Human Lung Microvascular Endothelial Cells (Lonza; Cat. #CC-2527) | 21TL169356 | Female | Caucasian | 2 | 34 | No | No | No | No | No |

**Supplementary Table 2.** Metadata for porcine donors of primary alveolar epithelial and primary lung microvascular endothelial cells.

| **Porcine donor** | **Cell type** | **Lot number** |
| --- | --- | --- |
| P1 | Porcine Primary Alveolar Epithelial Cells (Cell Biologics; Cat. #P-6053) | 061913NAB.JS |
| P2 | Porcine Primary Alveolar Epithelial Cells (Cell Biologics; Cat. #P-6053) | M051623W12 |
| P3 | Porcine Primary Lung Microvascular Endothelial Cells (Cell Biologics; Cat. #P-6011) | 012616.JS |

**Supplementary Table 3.** Antibodies used in immunofluorescence assays.

| Target Antigen | Antibody | Manufacturer/Provider | Cat. # | Organ-on-chip | Channel | Biosafety level |
| --- | --- | --- | --- | --- | --- | --- |
| **Primary antibodies** |  |  |  |  |  |  |
| Acetylated tubulin | Acetyl-α [acetylated] Tubulin (Lys40) (D20G3) XP<sup>®</sup> Rabbit mAb | Thermo Fisher Scientific | PA5-47488 | hsaOOC | Apical | 2 |
| CDH1 | Anti-E Cadherin antibody - Intercellular Junction Marker | Abcam | Ab15148 | hsaOOC | Apical | 2 |
| CDH5 | CD144 (VE-cadherin) Monoclonal Antibody (16B1), eBioscience | Thermo Fisher Scientific | 14-1449-82 | hsaOOC, paOOC | Basal | 2 |
| EFNB2 | Human/Mouse/Rat Ephrin-B2 polyclonal antibody | R&D Systems | AF496 | hsaOOC, paOOC | Apical and basal | 2 |
| MKI67 | Ki-67 Monoclonal Antibody (SolA15), FITC, eBioscience | Thermo Fisher Scientific | 11-5698-82 | hsaOOC, paOOC | Apical | 2 |
| MPO | Rabbit Myeloperoxidase Polyclonal Antibody | Thermo Fisher Scientific | PA516672 | hsaOOC | Apical and basal channels (in presence of TNF) | 4 |
| NiV G | Nipah virus glycoprotein G antibody, customized | Christopher Broder, Uniformed Health Services University | PA8905 | hsaOOC | Apical and basal | 4 |
| PECAM1 | CD31 Polyclonal Antibody | Thermo Fisher Scientific | PA5-32321 | hsaOOC, paOOC | Basal | 2 |
| TJP1 | ZO-1 Monoclonal Antibody (ZO1-1A12), Alexa Fluor 594 | Thermo Fisher Scientific | 339194 | hsaOOC, paOOC | Apical | 2 |
| Nucleic acid | Hoechst 33342, trihydrochloride trihydrate | Thermo Fisher Scientific | H3570 | hsaOOC, paOOC | Apical and basal | 2 |
| **Secondary antibodies** |  |  |  |  |  |  |
| / | Donkey anti-Goat IgG (H+L) Cross-Adsorbed Secondary Antibody, Alexa Fluor 647 | Thermo Fisher Scientific | A-21447 | hsaOOC | Apical | 2 |
| / | Donkey anti-Mouse IgG (H+L) Highly Cross-Adsorbed Secondary Antibody, Alexa Fluor Plus 594 | Thermo Fisher Scientific | A-32744 | hsaOOC | Apical | 2 |
| / | Donkey anti-Rabbit IgG (H+L) Highly Cross-Adsorbed Secondary Antibody, Alexa Fluor 488 | Thermo Fisher Scientific | A-21206 | hsaOOC | Apical | 2, 4 |
| / | Donkey anti-Rat IgG (H+L) Highly Cross-Adsorbed Secondary Antibody, Alexa Fluor 594 | Thermo Fisher Scientific | A-21209 | hsaOOC | Apical | 2 |

CDH1, cadherin 1; CDH5, cadherin 5; EFNB2, ephrin B2; hsaOOC, human small airway lung organ-on-chip; MKI67, marker of proliferation KI-67; MPO, myeloperoxidase; NiV G, Nipah virus glycoprotein; paOOC, porcine alveolus lung organ-on-chip; PECAM1, platelet and endothelial cell adhesion molecule 1; pro-SFTPB, pro-surfactant protein B; TNF, tumor necrosis factor; TJP1, tight junction protein 1; VWF, von Willebrand factor.

## Notes

### Competing Interest Statement

The authors have declared no competing interest.

